# Proteolytically activated antibacterial toxins inhibit the growth of diverse Gram-positive bacteria

**DOI:** 10.1101/2025.04.13.648598

**Authors:** Jake Colautti, Stephen R. Garrett, John C. Whitney

## Abstract

Many species of bacteria produce small-molecule antibiotics that enter and kill a wide range of competitor microbes. However, diffusible antibacterial proteins that share this broad-spectrum activity are not known to exist. Here, we report a family of proteins widespread in Gram-positive bacteria that display potent antibacterial activity against a diverse range of target organisms. Upon entering susceptible cells, these <u>a</u>nti<u>b</u>acterial <u>p</u>roteins (ABPs) enzymatically degrade essential cellular components including DNA, tRNA, and rRNA. Unlike previously characterized bactericidal proteins, which require a specific cell surface receptor and therefore display a narrow spectrum of activity, we find that ABPs act in a receptor-independent manner and consequently kill bacteria spanning multiple bacterial phyla. Target cell entry by ABPs requires proteolytic activation by a cognate, co-exported serine protease and the liberated toxin component of the cleaved ABP is driven across the target cell membrane by the proton motive force. By examining representative ABPs from diverse pathogenic, commensal, and environmental bacteria, we show that broad-spectrum antibacterial activity is a conserved property of this protein family. Collectively, our work demonstrates that secreted proteins can act as broad-spectrum antibiotics, suggesting that ABPs represent one of potentially many such families produced in nature.

**Significance Statement:** Many bacteria produce proteins with antibacterial properties. However, owing to their reliance on a specific surface receptor for target cell entry, all known antibacterial proteins are only active against a narrow range of organisms. Using biochemical and genetic approaches, this study reports the discovery of a new family of antibacterial proteins secreted by many Gram-positive bacteria that enter and kill a broad spectrum of bacteria. Entry of these proteins into susceptible bacteria does not require a receptor and instead relies on cleavage by a co-secreted protease and the proton motive force of the target cell. Overall, our findings reveal a new family of antibacterial proteins and provides insight into how these proteins enter and kill a broad range of bacteria.

## Introduction

Antibiotics synthesized by bacteria act by diffusing away from the producing cell and inhibiting the growth of nearby competitor species (1, 2). In most cases, these secreted molecules can enter and kill diverse species of bacteria, which has led to their extensive use in medicine and agriculture (3, 4). In addition to small-molecule antibiotics, some bacteria also export proteins with antibacterial properties (5, 6). These proteins typically function as toxic enzymes that kill susceptible bacteria by disrupting essential cellular pathways (7, 8). In most instances, delivery of these large macromolecules into target bacteria relies on direct injection by sophisticated protein secretion machines that require prolonged cell-to-cell contact to facilitate interbacterial killing (7, 9–15). However, like small-molecule antibiotics, some antibacterial proteins diffuse through the environment and spontaneously enter susceptible cells. The best described among these proteins are colicins, which mediate competition between closely related strains of *Escherichia coli* (6, 16). Colicins enter competitor cells by binding to and translocating through cell surface proteins (17–19). Due to their reliance on highly specific protein-protein interactions for target cell entry, all known diffusible antibacterial proteins, including colicins, have a narrow spectrum of activity (6, 20). To date, no secreted protein has been described that inhibits the growth of a wide range of distantly related bacteria.

## Results

### A secreted protein with broad-spectrum antibiotic activity

A recent bioinformatic analysis of bacterial genomes identified proteins encoded by diverse Gram-positive bacteria that share homology with the antibacterial toxin domains of several well-characterized colicins (Figure 1A)(21). In addition to a C-terminal antibacterial toxin domain, these proteins contain an N-terminal domain of unknown function, termed BetaH, and a signal peptide that targets them for co-translational export through the general secretory pathway (Sec). Furthermore, these proteins invariably co-occur with a predicted serine protease, which also contains an N-terminal Sec signal peptide and is therefore predicted to be co-secreted alongside its associated antibacterial protein. Lastly, these putative antibacterial proteins are encoded upstream of genes that are expected to confer immunity to their toxic activity, thus protecting producing organisms (21).

**Figure 1:**
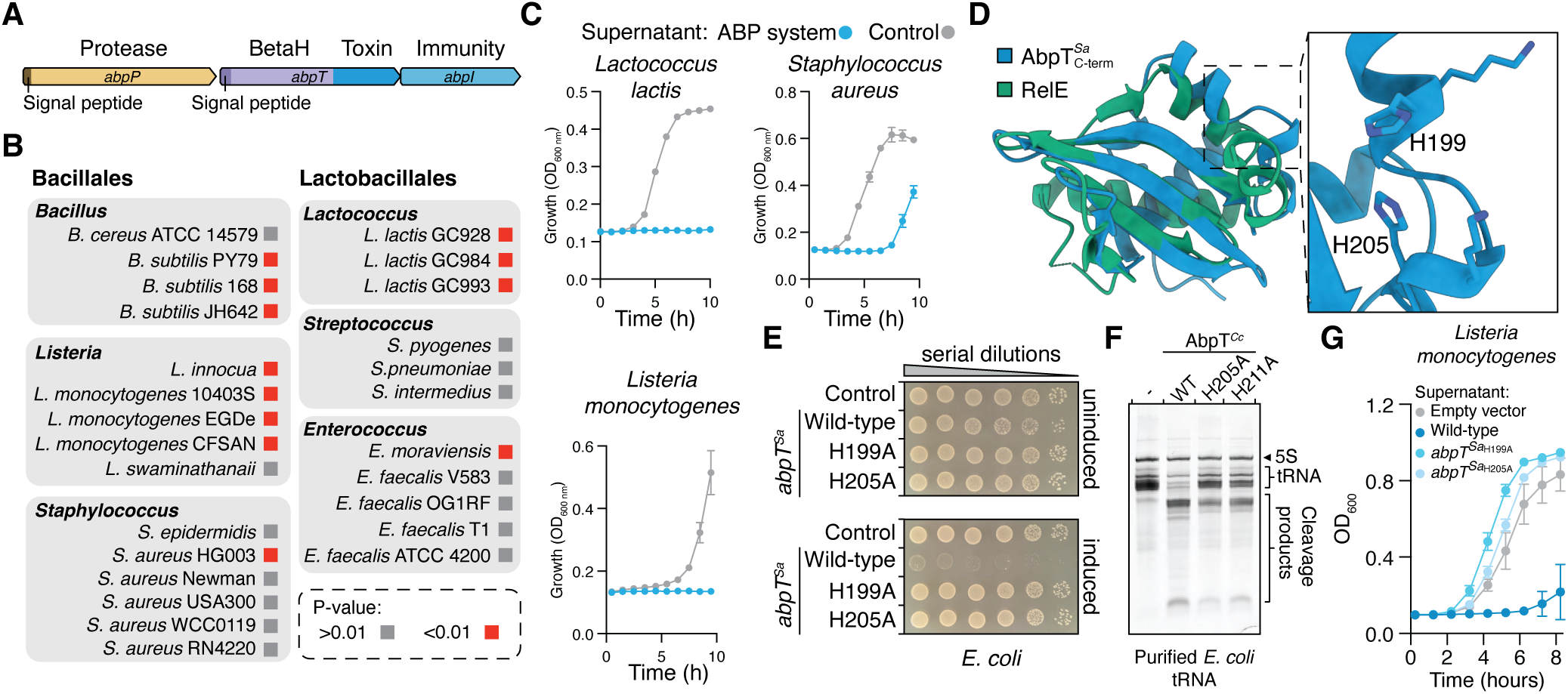
An ABP gene cluster encodes a secreted protein with broad-spectrum antibiotic activity. A) Schematic representation of an ABP gene cluster encoded by a human clinical isolate of *Staphylococcus aureus* (NCBI BioSample: SAMN38329808) B) Representative ABP susceptibility screening results. P values were calculated from a one-tailed homoscedastic T test comparing growth of the indicated bacterial strain grown in the presence or absence of supernatant from ABP-expressing *S. aureus*. Screening results of additional bacterial strains are presented in supplemental Figure 1. C) Growth of the indicated bacterial species treated with supernatant from *S. aureus* expressing the ABP gene cluster depicted in (A). D) AlphaFold3 model of the C-terminal BECR domain of AbpT*^Sa^* overlaid with the RelE ribonuclease (PDB 3BPQ). The catalytic histidine dyad of AbpT*^Sa^* is highlighted in the inset. E) Viability of *E. coli* harbouring plasmids encoding *abpT^Sa^* or the indicated *abpT^Sa^* variants grown on solid medium in the presence or absence of inducer, as indicated. F) Urea PAGE analysis of purified *E. coli* tRNA treated with purified AbpT*^Cc^* or the indicated AbpT*^Cc^* variants. Dashed line denotes RNA treated with buffer only. RNA was visualized by staining with SYBR Gold. G) Growth of *Listeria monocytogenes* treated with concentrated supernatant from *S. aureus* expressing the ABP gene cluster depicted in (A) or the indicated ABP gene cluster variants. In panels C and G, data are presented as mean +/- SEM, n=3 biological replicates.

Given that secreted antibacterial proteins have been extensively characterized in Gram-negative bacteria but remain elusive among Gram-positive organisms, we reasoned that these genes may encode secreted toxins that are functionally analogous to colicins and thus would inhibit the growth of a narrow spectrum of Gram-positive targets. To identify susceptible organisms, we screened supernatant from a culture of *Staphylococcus aureus* expressing a candidate gene cluster for growth inhibitory activity against a diverse panel of Gram-positive and Gram-negative bacteria. Strikingly, this culture supernatant not only inhibited the growth of related *S. aureus* strains but also of many distantly related Gram-positive bacteria belonging to the orders Bacillales and Lactobacillales (Figure 1B,C and Figure S1A). This finding indicates that although this protein displays antimicrobial activity as predicted, its antibiotic spectrum is much broader than colicins or other previously described diffusible antibacterial proteins. To reflect this newfound activity, we henceforth refer to these tricistronic gene clusters as <u>a</u>nti<u>b</u>acterial <u>p</u>rotein systems (ABPs), which are defined by genes encoding a serine <u>p</u>rotease (*abpP*), an antibacterial toxin protein (*abpT*), and an immunity protein *(abpI*).

To gain further insight into the mechanisms by which ABPs inhibit bacterial growth, we first sought to functionally characterize the C-terminal toxin domain of *S. aureus* AbpT (AbpT*^Sa^*). This domain is predicted by AlphaFold3 to adopt the Barnase/EndoU/Colicin D/RelE (BECR) fold shared by many ribonucleases (Figure 1D)(22–24). Furthermore, AbpT*^Sa^* contains a dyad of conserved histidine residues that are frequently present in the active site of metal-independent ribonucleases (8, 25). Consistent with this predicted biochemical activity, expression of AbpT*^Sa^* in the cytoplasm of *Escherichia coli* inhibits growth and causes rapid RNA degradation (Figure 1E, Figure S1B,C). Recombinantly produced AbpT*^Sa^* was unstable in solution, but a homologous AbpT encoded by *Corynebacterium casei* (AbpT*^Cc^*; 33.7% BECR domain seq. ID) cleaved multiple RNA species *in vitro*, mirroring the ribonuclease activity we observe in cells (Figure 1F). Mutagenesis of the AbpT*^Sa^* catalytic dyad abrogates the growth inhibitory activity of our *S. aureus* supernatant, indicating that its antibacterial properties arise from secreted AbpT’s ribonuclease activity (Figure 1G). Finally, co-expression of AbpI*^Sa^* in cells expressing cytoplasmic AbpT*^Sa^*confers protection against the antibacterial activity of AbpT*^Sa^*, supporting our hypothesis that AbpI proteins function as AbpT-specific immunity proteins (Figure S1D). Together, these results indicate that AbpT*^Sa^* is an antibacterial ribonuclease that inhibits the growth of a diverse range of bacteria and its activity is neutralized by a cognate immunity protein, AbpI*^Sa^*.

### Cognate serine proteases activate antibacterial AbpT proteins

The strict co-occurrence between AbpT proteins and an adjacently encoded serine protease suggests that these proteases contribute to the function of ABP systems. Indeed, supernatant from a *S. aureus* strain expressing a catalytically inactive variant of its AbpP protease displays attenuated antibacterial activity against *L. monocytogenes* (Figure 2A). AbpP belongs to the S8 family of serine proteases, which are involved in the proteolytic cleavage of inactive pro-proteins into their active forms in diverse biological contexts (21, 26, 27). We therefore hypothesized that AbpP activates its associated antibacterial AbpT protein by proteolytic cleavage. Consistent with this idea, we only detect a smaller fragment of AbpT*^Sa^*in supernatant from a *S. aureus* culture expressing the wild-type ABP*^Sa^*system whereas supernatant from culture expressing the catalytically inactive AbpP*^Sa^* protease contains unprocessed AbpT*^Sa^*at its expected molecular weight (Figure 2B). Given that AbpP and AbpT both possess Sec signal sequences, we conclude that AbpP proteases cleave their co-exported antibacterial proteins following their secretion from producing cells.

**Figure 2:**
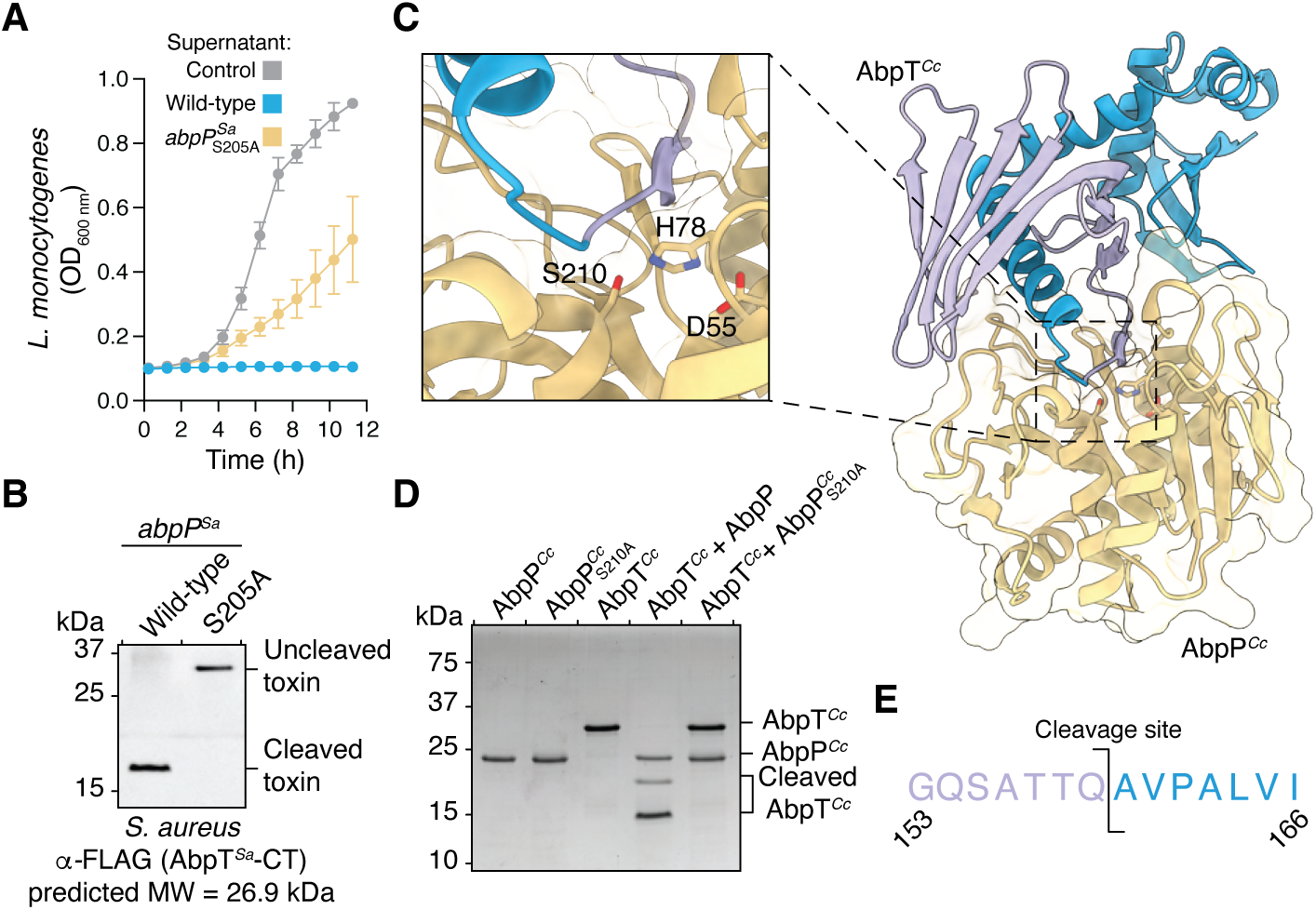
Cleavage by a cognate serine protease activates ABP toxins. A) Growth of *L. monocytogenes* treated with supernatant from *S. aureus* expressing the wild-type ABP gene cluster or a mutant gene cluster in which the predicted catalytic serine of AbpP has been substituted to alanine (S205A). Supernatant from *S. aureus* harbouring the empty vector is used as a control. Data are presented as mean +/- SEM. B) Western blot analysis of supernatant from *S. aureus* cultures in panel A. In both strains, a FLAG epitope tag was fused to the C-terminus of AbpT*^Sa^* to facilitate protein detection. C) AlphaFold3 model of the predicted complex formed by AbpT*^Cc^*and AbpP*^Cc^*. The predicted N- and C-terminal cleavage fragments of AbpT*^Cc^* by the indicated catalytic triad of AbpP*^Cc^* are coloured purple and blue, respectively. AbpP*^Cc^* is depicted in orange cartoon representation. D) AbpP*^Cc^* cleaves AbpT*^Cc^* into two distinct fragments. SDS-PAGE analysis of the indicated proteins or protein mixtures, visualized by staining with Coomassie Brilliant Blue. E) Schematic representation of the experimentally determined AbpP*^Cc^* cleavage site in AbpT*^Cc^* depicting the sequence at the N-terminal (purple) and C-terminal (blue) sides of the cleavage site. N-terminal sequencing data are available in supplemental file S1.

To gain further insight into the consequences of AbpT cleavage by AbpP proteases, we predicted the structures of these proteins using AlphaFold3. The conserved N-terminal BetaH domain of AbpT contains a curved seven-stranded β-sheet joined to a central α-helix by a short unstructured loop (Figure S2A) (21). AlphaFold3 generates a highly confident prediction of the complex formed between AbpT and AbpP in which this unstructured loop is positioned within the protease active site (Figure S2B). This prediction suggests that AbpP proteases cleave AbpT antibacterial proteins between the β-sheet of the BetaH domain and its central α-helix, which would produce a C-terminal ribonuclease domain-containing fragment that is consistent with the molecular weight of the fragment present in our *S. aureus* culture supernatant (Figure 2B and Figure S2B). To experimentally test the predicted protease cleavage site, we attempted to reconstitute AbpP cleavage of AbpT *in vitro* using the *C. casei* ABP system. As is the case for the *S. aureus* ABP system, AlphaFold3 confidently positions the unstructured loop of AbpT*^Cc^*in the active site of AbpP*^Cc^* (Figure 2C and Figure S2C). Consistent with this model and our observations in *S. aureus* culture supernatants, purified AbpP*^Cc^* cleaves AbpT*^Cc^* into two distinct fragments in a manner that is dependent on its catalytic serine residue (Figure 2D). N-terminal sequencing of the resulting AbpT*^Cc^* fragments revealed that cleavage occurs between Gln159 and Ala160, which is the exact site predicted by the AlphaFold3 model of the complex (Figure 2E). Finally, we find that proteolytic cleavage only occurs between cognate AbpP-AbpT pairs, strongly suggesting that cleavage at this site is the evolved function of these serine proteases (Figure S3A and B).

### Translocation of antibacterial proteins into target cells

Given the tricistronic arrangement of ABP systems, we reasoned that the co-exported antibacterial protein and its cognate protease likely constitute the minimal functional unit needed for bacterial growth inhibition. To investigate this possibility, we examined the antibiotic properties of purified AbpT protein following cleavage by its associated AbpP protease. Strikingly, we found that the addition of proteolytically cleaved AbpT*^Cc^*to bacterial lawns of diverse Gram-positive organisms resulted in a clear zone of killing, indicating that cleaved AbpT proteins do not require any additional cellular factors for their antibacterial activity (Figure 3A). Consistent with our findings using *S. aureus* supernatants, uncleaved AbpT*^Cc^* displays substantially attenuated but still detectable growth inhibitory activity (Figure 3A). Treatment of susceptible bacteria with purified AbpT*^Cc^* protein also leads to the intracellular accumulation of RNA degradation fragments that mirror the fragments produced by AbpT*^Cc^ in vitro*, indicating that the RNAse activity of AbpT*^Cc^* is responsible for its antibacterial properties (Figure 3B). This finding is further confirmed by the observation that site-specific mutagenesis of the catalytic histidine dyad in the ribonuclease domain of AbpT*^Cc^* completely attenuates its antibacterial activity (Figure C and Figure S4A). To quantify the contribution of proteolytic cleavage to the antibacterial activity of AbpT, we next determined the minimum inhibitory concentration (MIC) of both cleaved and uncleaved AbpT*^Cc^* against a panel of diverse bacteria and found that AbpP*^Cc^*cleavage enhances AbpT*^Cc^*activity by approximately 20-fold (Figure 3D and Figure S4B-D). Furthermore, like many small-molecule antibiotics, we find that cleaved AbpT*^Cc^* is bactericidal, rather than bacteriostatic, at its MIC (Figure S4E-G). Based on these findings, we conclude that proteolytic cleavage by AbpP strongly enhances the bactericidal activity of AbpT proteins and that together these two proteins constitute the minimal functional unit of ABP systems.

**Figure 3:**
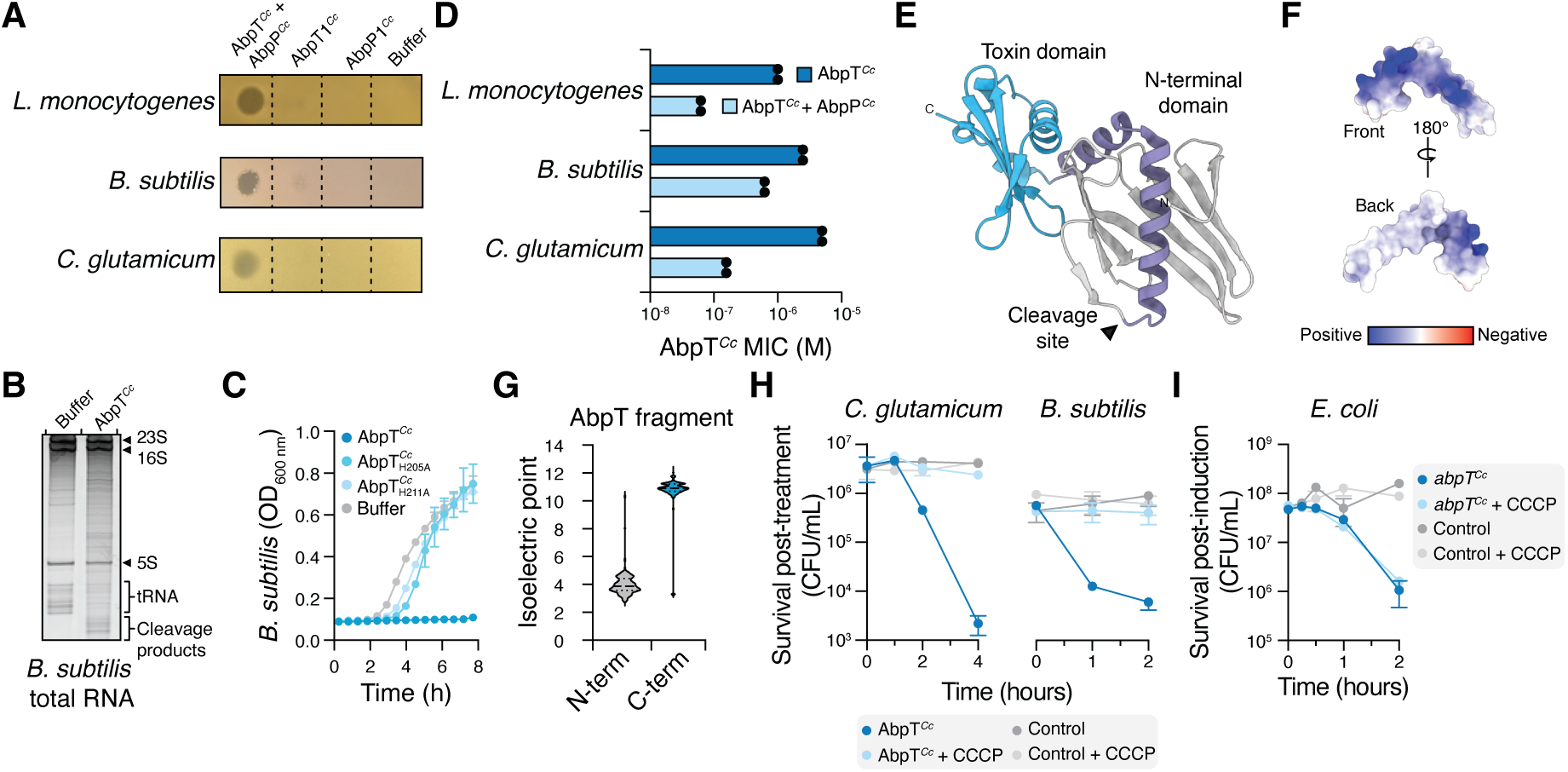
ABP toxins require the proton motive force for entry into the bacterial cytoplasm A) Agar diffusion assays of the indicated purified proteins spotted onto diverse Gram-positive bacteria. B) Urea PAGE analysis of total RNA extracted from *B. subtilis* treated with proteolytically cleaved AbpT*^Cc^* or buffer control. RNA was visualized by staining with SYBR Gold. C) Growth of *B. subtilis* treated with purified proteolytically cleaved AbpT*^Cc^* or the indicated single amino acid substitutions. D) Minimum inhibitory concentration (MIC) assays of purified AbpT*^Cc^*alone or in combination with equimolar AbpP*^Cc^*. Complete MIC assay results are presented in supplemental figure 4B-D. E) AlphaFold3 model of AbpT*^Cc^* depicting the N-terminal BetaH domain (grey and purple) and C-terminal toxin domain (blue). The AbpP*^Cc^* cleavage site is indicated by the arrow. F) Surface electrostatic representation of the central α-helix liberated by AbpP*^Cc^* cleavage (shown as purple in panel E). G) Violin plot of the predicted isoelectric points of the N- and C-terminal proteolytic fragments of 503 representative ABP toxins. The mean predicted isoelectric points of the N- and C-terminal fragments are 4.1 and 10.7, respectively. H) Survival of the indicated bacterial species following incubation with exogenously added proteolytically cleaved AbpT*^Cc^* in the presence or absence of the membrane-depolarizing agent CCCP. Control represents cultures treated with buffer. I) Survival of *E. coli* expressing *abpT^Cc^* or a control strain harbouring the empty vector in the presence or absence of the membrane-depolarizing agent CCCP. In panels C, H, and I, data are presented as mean +/- SEM, n=3 biological replicates.

Due to their reliance on specific interactions with a receptor protein, known diffusible antibacterial proteins display an extremely narrow spectrum of activity, typically restricted to a subset of strains of a single organism (6, 28). The observation that AbpT proteins enter and kill bacteria belonging to multiple phyla coupled with their lack of a receptor binding domain suggests that these protein antibiotics do not require a receptor to enter susceptible cells. Therefore, we next sought to understand how these large proteins access the bacterial cytoplasm without the involvement of a protein receptor. Given our data showing that the proteolytic separation of the C-terminal ribonuclease domain and central α-helix from the N-terminal β-sheet activates AbpT, we reasoned that the physical properties of the central α-helix and/or ribonuclease domain must contribute to the translocation of cleaved AbpT into the cytoplasm of susceptible cells. Interestingly, the central α-helix in AbpT proteins is amphipathic, and its hydrophobic face is shielded from the aqueous milieu by its N-terminal β-sheet prior to cleavage by AbpP (Figure S3E and F). Additionally, the C-terminal toxin domains of AbpT proteins are highly cationic, with an average isoelectric point of 10.2, despite their predicted enzymatic diversity (Figure 3G and Figure S5A). Amphipathic α-helices and basic amino acids enable proteins to spontaneously enter lipid bilayers and are therefore frequently present in membrane active proteins such as cationic antimicrobial peptides (6, 29, 30). Therefore, the conserved presence of these two properties in cleaved AbpT proteins led us to speculate that they spontaneously insert into the cytoplasmic membrane and are driven across this membrane by the proton motive force (PMF), which is known to provide the energy required for membrane translocation of cationic antimicrobial peptides and cationic small-molecule antibiotics (6, 31–33). Consistent with this hypothesis, chemical dissipation of the PMF using <u>c</u>arbonyl <u>c</u>yanide-m-<u>c</u>hloro<u>p</u>henylhydrazone (CCCP) protects distantly related bacteria from the bactericidal activity of exogenously added purified AbpT*^Cc^* (Figure 3H). By contrast, dissipation of the PMF does not provide protection from intracellular AbpT*^Cc^* expression (Figure 3I). Together, these findings indicate that the PMF is not required for the lethal ribonuclease activity of AbpT*^Cc^* but is essential for the protein’s ability to access the bacterial cytoplasm.

### ABPs are widespread in bacteria and encode functionally diverse protein antibiotics

Despite belonging to different bacterial phyla, the observation that *S. aureus* and *C. casei* encode AbpTs with the same mechanism of action suggests that ABPs in general function as broad-spectrum protein antibiotics. To explore this hypothesis further, we next set out to characterize the spectrum of activity and mode of action of ABP systems from diverse Gram-positive bacteria. To identify candidate ABP systems, we constructed a phylogenetic tree of AbpT proteins using their conserved N-terminal BetaH domains (Figure 4A). Encouragingly, the BetaH domains present in the AbpTs of *S. aureus* and *C. casei* map to distinct but adjacent clades of this tree, suggesting that their broad-spectrum antibiotic activity is not restricted to a subset of ABPs but rather is a general feature of this protein family (Figure 4A). To bolster this assertion, we selected a more distantly related AbpT homolog encoded by *Priestia megaterium* (AbpT*^Pm^*) for further characterization. Despite having a C-terminal toxin domain that is unrelated to that of AbpT*^Sa^*or AbpT*^Cc^*, AbpT*^Pm^* is also predicted to adopt the BECR ribonuclease fold (Figure 4B)(34). After confirming AbpT*^Pm^* expression inhibits bacterial growth and that AbpI*^Pm^* co-expression alleviates this growth inhibitory activity, we examined the biochemical activity of purified AbpT*^Pm^* (Figure 4C and Figure S6A). In contrast to the closely related AbpT*^Sa^* or AbpT*^Cc^*proteins, we found that AbpT*^Pm^*displays highly promiscuous ribonuclease activity *in vitro* and thus acts by a distinct mechanism of action (Figure 4D) (35). Despite this difference in protein function, AbpT*^Pm^* displays a comparable antibiotic spectrum of activity, confirming that it too functions as a broad-spectrum protein antibiotic (Figure 4E). Based on these findings, we conclude that broad spectrum activity is a general feature of ABP systems.

**Figure 4:**
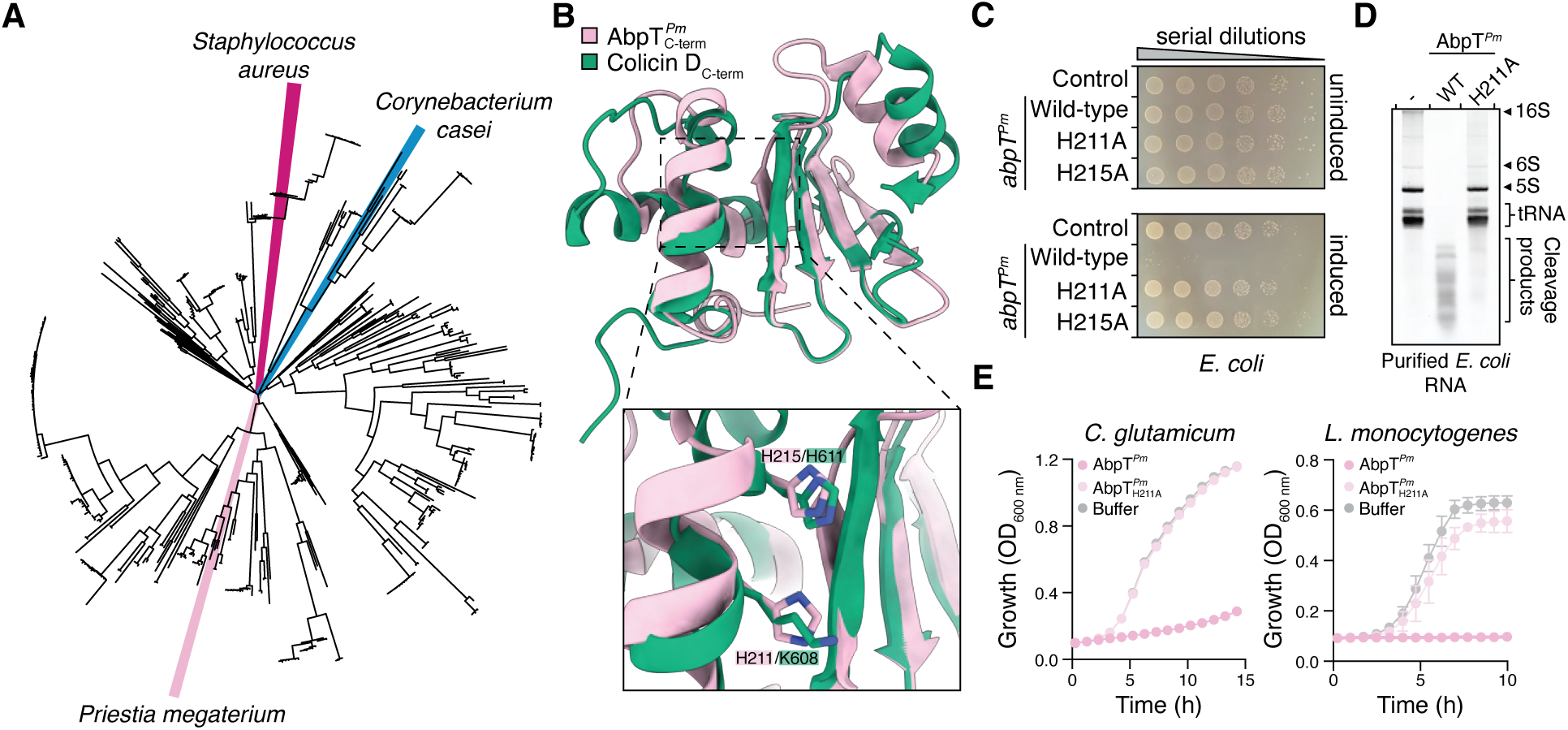
Functionally diverse ABP systems encode broad-spectrum protein antibiotics. A) Phylogenetic tree of the N-terminal BetaH domain of AbpT. AbpT proteins characterized in this study are indicated by coloured wedges. B) AlphaFold3 model of the C-terminal toxin domain of AbpT*^Pm^* overlaid with the C-terminal toxin domain of colicin D (PDB 1V74). Active site residues of both proteins are presented in the inset. C) Viability of *E. coli* harbouring plasmids encoding wild-type AbpT*^Pm^* or the indicated AbpT*^Pm^* variants grown on solid media in the presence or absence of inducer, as indicated. D) Urea PAGE analysis of purified *E. coli* RNA treated with purified AbpT*^Pm^*or a purified AbpT*^Pm^*catalytic mutant. Dashed line denotes RNA treated with buffer only. Total RNA was visualized by staining with SYBR Gold. E) Growth of the indicated bacterial species treated with purified AbpT*^Pm^*, the indicated single amino acid substitution, or buffer only. Data are presented as mean +/- SEM, n=3 biological replicates.

We next examined the toxin domains present in the ABP systems identified by our bioinformatic search to better understand the diversity of mechanisms by which these proteins kill susceptible bacteria. In some instances, these C-terminal toxin domains share structural similarity to toxins delivered between bacterial cells using cell contact-dependent interbacterial antagonism pathways (21). For example, the epidemic methicillin-resistant *S. aureus* strain MRSA252 encodes an AbpT protein with a toxin domain that is homologous to CdiA^ECL^, a ribosome-targeting toxin produced by a <u>C</u>ontact-<u>D</u>ependent Inhibition (CDI) pathway in *Enterobacter cloacae* (Figure S7A and B) (36, 37). We experimentally validated that this AbpT shares the same biochemical activity as CdiA^ECL^ by showing that its expression inhibits bacterial growth in a manner dependent on a catalytic residue conserved with CdiA^ECL^, and that growth-inhibited cells undergo cleavage at the 3’ end of their 16S ribosomal RNA (Figure S7C-F)(36). Other ABP toxins share striking structural homology to restriction endonucleases or HNH nucleases, both of which act as metal-dependent DNAses (Figure S8A, 9A) (25, 38). Expression of these predicted nucleases inhibits the growth of *E. coli* and causes degradation of the bacterial chromosome (Figure S8B-E, S9B-E). In addition to C-terminal domains that share homology with previously described antibacterial toxins, many other ABP systems encode AbpT proteins that do not share sequence or structural similarity to proteins of known function and therefore likely represent antibacterial proteins with novel modes of action (Figure S10A,B) (21). For example, expression of one such AbpT encoded by *Bacillus cereus* (AbpT*^Bc^*) potently inhibits bacterial growth but its sequence and predicted structure offer no insight into the mechanism of its antibacterial activity (Figure S10C-F). Taken together, our results indicate that ABP systems encode functionally diverse broad-spectrum protein antibiotics that are widespread among Gram-positive bacteria.

## Discussion

We have discovered that ABPs are a large family of secreted bacterial proteins that function as broad-spectrum antibiotics. These proteins are activated by a cognate protease and kill susceptible cells by multiple distinct mechanisms that target conserved essential processes in bacteria. Unlike other antibacterial proteins that rely on a specific receptor for entry into susceptible cells, ABPs are spontaneously driven into the cytoplasm of a wide range of bacteria using the proton motive force. The finding that ABP systems are present in diverse Gram-positive organisms suggests that these proteins are abundant in nature. Together, these findings establish that it is possible for diffusible proteins to act intercellularly to inhibit the growth of a broad spectrum of bacterial competitors.

Although many families of diffusible antimicrobial proteins have been identified, none appear comparable to those encoded by ABP systems (6, 20, 39). ABPs have some similarities to colicins; both families of proteins act intercellularly to disrupt bacterial physiology, and, in many instances, use the same enzymatic toxin domains. However, ABPs are both produced by and are active against a broad spectrum of Gram-positive bacteria whereas colicins are produced by *E. coli* and are only active against a narrow subset of strains within this Gram-negative species (6). In this sense, ABPs are perhaps more appropriately comparable to <u>a</u>nti<u>m</u>icrobial <u>p</u>eptides (AMPs), which are secreted proteinaceous molecules produced by Gram-positive bacteria that inhibit the growth of a wide range of non-self Gram-positive competitors (40–42). Furthermore, the presence of an outer membrane protects Gram-negative bacteria against the action of most AMPs and likely also explains the resistance of these organisms to ABPs. However, AMPs are not multi-domain enzymes but rather short peptides that frequently act by disrupting cell envelope integrity (40, 43). Therefore, the ABP systems that we report herein appear to combine the enzymatic properties of colicins with the broad-spectrum, receptor-independent activity of AMPs.

Our discovery of ABP systems raises many important questions for future investigation. The finding that ABPs are widespread in Gram-positive bacteria suggests that these proteins contribute to bacterial competition. However, the environmental conditions under which these systems mediate such antagonistic interbacterial interactions remains unknown. Here, we focus on ABP systems encoded by *Corynebacterium* and *Staphylococcus* species, which are known to compete for occupancy within the human skin and nasal microbiomes (44, 45). The observation that these organisms both encode and are susceptible to ABPs suggests that these systems contribute to competition within these human niches. For instance, ABP systems could contribute to the pathogenicity of some *S. aureus* strains by enabling these strains to outcompete commensal *Corynebacterium* species. Consistent with this possibility, an AbpT protein is detectable in the secreted proteome of a *S. aureus* clinical isolate, suggesting that production of an ABP might contribute to the pathogenicity of this strain (46). Alternatively, ABP systems encoded by commensal *Staphylococcus* and/or *Corynebacterium* species may allow an established bacterial community to prevent host colonization by an ABP-susceptible pathogen. Additionally, ABP systems are present in numerous plant-associated bacteria that are known to adopt commensal or pathogenic lifestyles (47, 48). The extent to which ABP systems contribute to bacterial competition in plant-associated niches therefore present additional opportunities for further study. Lastly, the finding that it is possible for large proteins to act as broad-spectrum antibiotics suggests that there may exist other secreted protein families with this activity. It will also be pertinent to consider potential biotechnological or therapeutic applications of ABPs because they represent a diverse class of functionally novel protein antibiotics. In summary, our work demonstrates that large enzymatic proteins can function as broad-spectrum antibiotics, offers new opportunities to expand our understanding of bacterial antagonism, and lays the groundwork for the development of new classes of protein antibiotics for clinical or industrial use.

## Materials and Methods

### Bacterial strains and growth conditions

A complete list of strains and growth conditions used in this study can be found in supplementary table 1, and all strains are available upon request from the corresponding authors. *Escherichia coli* strain XL1 Blue (Novagen) and DC10b were used for plasmid maintenance and strain BL21 (DE3) pLysS (Novagen) was used for recombinant protein expression. *E. coli* strains were grown in lysogen broth (LB) medium (10 g/L tryptone, 5 g/L yeast extract, 10 g/L NaCl) at 37°C shaking at 220 revolutions per minute (RPM). *E. coli* cultures were supplemented with 50 μg/mL kanamycin, 100 μg/mL ampicillin, 25 μg/mL chloramphenicol, 200 μg/mL trimethoprim, 15 μg/mL gentamicin, 0.1% L-rhamnose, 0.1% L-arabinose, and 1 mM isopropyl β-D-1-thiogalactopyranoside, as appropriate. *Staphylococcus aureus* cultures were grown in tryptic soy broth (TSB) medium (Bacto) at 37°C shaking at 220 revolutions per minute (RPM). *S. aureus* cultures were supplemented with 10 μg/mL chloramphenicol and 500 ng/mL anhydrotetracycline as appropriate.

### DNA manipulation and plasmid construction

All expression vectors were constructed using standard restriction enzyme-based cloning procedures (49). Primers were synthesized by Integrated DNA Technologies. Phusion polymerase, restriction enzymes, and T4 DNA ligase were obtained from New England Biolabs. Whole plasmid sequencing was performed by Plasmidsaurus using Oxford Nanopore Technology with custom analysis and annotation. Oligonucleotide sequences and plasmids used in this study are provided in supplementary tables 2 and 3, respectively.

### Preparation of concentrated supernatant for bacterial growth inhibition assays

*S. aureus* strains derived from the sequenced strain USA300 harbouring the indicated pRAB11 vectors were grown overnight at 37°C shaking at 220 RPM in 5 mL TSB supplemented with 10 μg/mL chloramphenicol. Overnight cultures were diluted to OD600 of 0.1 in 500 mL TSB with 10 μg/mL chloramphenicol and grown at 37°C shaking at 220 RPM to an OD600 of 0.5. Anhydrotetracycline was added to a final concentration of 500 ng/mL to induce gene expression. Upon reaching OD600 of 8.0, cultures were pelleted at 6000 *g* for 20 minutes and protein was precipitated from culture supernatant by the addition of ammonium sulfate to a final concentration of 80% (w/v). Supernatant was stirred overnight at 4°C and subsequently centrifuged at 36 000 *g* for 45 minutes. Precipitated supernatant was resuspended in 10 mL 1x phosphate buffered saline (PBS) pH 7.4 and dialyzed twice against 2 L 1x PBS at room temperature for 1 hour each. Dialyzed protein was further concentrated using a 10 kDa molecular weight cutoff centrifugal filter device (MilliporeSigma) to a final volume of 2 mL (250-fold concentration of original culture supernatant). Concentrated protein was snap frozen in liquid nitrogen and stored at −80°C before use.

### Bacterial growth inhibition screening

Strains used in this assay, their sources, and growth conditions are listed in supplementary table 1. Starting cultures were diluted to an OD of 0.05 in 90 μL of appropriate media in adjacent wells of a 96 well plate. 10 μL of concentrated culture supernatant (see above) collected from *S. aureus* cultures expressing either the indicated ABP system or an empty vector control was added to each well. Plates were subsequently incubated in growth conditions appropriate for each strain for 8 hours, and OD600 was measured using a BioTek Synergy H1 multimode reader (Agilent). Two biological replicates of the screen were conducted, and the p-values were calculated using an unpaired T-test comparing mean OD600 of the culture treated with ABP supernatant compared with the culture treated with control supernatant.

### Bacterial growth curves

To validate initial hits from the ABP susceptibility screen, stationary phase cultures of the indicated bacterial strains grown in appropriate media were subcultured 1:100 into 96 well plates containing 90 μL of appropriate media, to which 10 μL concentrated ABP or control culture supernatant was added. Cultures were grown at the indicated temperatures in the presence or absence of shaking in a BioTek Synergy H1 multimode reader (Agilent) and the absorbance at 600 nm was measured every 30 minutes.

To determine the growth inhibition activity of purified AbpT proteins, experiments were conducted as above, but the indicated purified proteins were added at 1.5x their MIC instead of concentrated culture supernatant.

### E. coli toxicity experiments

Cultures of *E. coli* XL1 Blue harbouring the indicated plasmids were grown overnight in 2 mL LB supplemented with appropriate antibiotics at 37°C shaking at 220 RPM. Overnight cultures were normalized to an OD600 of 1.0 and were diluted in 10-fold series in a 96 well microtiter plate. 6 μL of 10^-1^-10^-6^ dilutions were spotted on LB 1.5% (w/v) agar containing 0.1% (w/v) L-rhamnose or 0.1% (w/v) L-arabinose to induce toxin expression from pSCrhaB2-CV- or pBAD33-derived vectors, respectively, or 1 mM IPTG to induce immunity expression from pPSV39-derived vectors. Plates were incubated at 37°C for 18 hours and photographed.

### RNA extraction

Cultures of *E. coli* XL1 Blue harbouring the indicated plasmids were grown overnight in 2 mL LB supplemented with appropriate antibiotics at 37°C shaking at 220 RPM. Cultures were diluted to OD600 0.1 in 25 mL LB supplemented with appropriate antibiotics and grown at 37°C shaking at 220 RPM for approximately 2 hours to OD600 of 0.5. Toxin expression was induced by the addition of L-rhamnose to a final concentration of 0.1% (w/v). After the indicated timepoints had elapsed, 5 mL culture was collected by centrifugation at 4000 *g* for 10 minutes at room temperature and the pellet was resuspended in 1 mL TRIzol reagent (ThermoFisher). Total RNA was extracted using a standard protocol (50). Briefly, *E. coli* resuspended in TRIzol reagent was vortexed for 30 seconds and incubated at room temperature for 5 minutes before the addition of 200 μL chloroform. The mixture was vortexed for 30 seconds and incubated on ice for 2 minutes before centrifugation at 14 000 *g* for 15 minutes at 4°C. 500 μL of the upper aqueous phase was pipetted into 500 μL of ice cold isopropanol, vortexed for 30 seconds, incubated on ice for 20 minutes, and centrifuged at 21 000 *g* for 15 minutes at 4°C. The resulting RNA pellet was washed three times with 1 mL ice cold 75% (v/v) ethanol and dried under a vacuum. The RNA pellet was resuspended in 100 μL 1x DNAse I buffer (NEB) to which 4 units RNAse-free DNAse I (NEB) was added. The reaction was incubated for 30 minutes at 37°C and RNA was subsequently extracted using a Monarch Spin RNA cleanup kit (NEB) following the manufacturer’s instructions. RNA was quantified using a NanoDrop spectrophotometer (ThermoFisher) and 500 ng of total RNA was analyzed by UREA PAGE stained with SYBR Gold (ThermoFisher).

The method above was used to extract RNA from *Bacillus subtilis* treated with purified AbpT, with the following modifications. An isolated colony of *B. subtilis* strain PY79 was collected from a LB plate containing 1.5% (w/v) agar and grown to OD600 of 0.5 in 10 mL LB at 37°C shaking at 220 RPM. The 10 mL culture was subsequently split into 2x 5 mL cultures; one culture was treated with purified AbpT*^Cc^* and AbpP*^Cc^* to a final concentration of 1 μM each and the other 5 mL culture was treated with an equal volume of sterile buffer (150 mM NaCl, 20 mM HEPES NaOH pH 7.5). Cultures were subsequently incubated at 37°C shaking at 220 RPM for 1 h before cells were collected by centrifugation at 4000 *g* at room temperature for 10 minutes. Pellets were washed 3x with 1 mL 1x PBS pH 7.4. Pellets were resuspended in 200 μL 1x PBS pH 7.4 containing 50 mg/mL hen egg lysozyme and incubated for 20 minutes at 37°C. 1 mL TRIzol reagent was subsequently added, and RNA extraction was completed as described above.

### RNAse activity assays

To determine the ribonuclease activity of AbpT*^Cc^* and AbpT*^Pm^ in vitro*, 1 μM of the indicated purified proteins was incubated with 2000 ng of total *E. coli* RNA extracted from *E. coli* strain MG1655 using the method described above or 2000 ng purified *E. coli* tRNA (Roche) in a 6 μL reaction volume in buffer containing 150 mM NaCl, 20 mM HEPES NaOH pH 7.5. Reactions were allowed to proceed for 20 minutes at 37°C and were quenched by the addition of an equal volume of 2x RNA loading dye (95% v/v formamide, 0.025% bromophenol blue, 0.025% SDS, 10 mM EDTA pH 8.0). Samples were subsequently heated to 95°C for 10 minutes and rapidly cooled by submerging in ice water. 500 ng total RNA was analyzed by UREA PAGE stained with SYBR Gold (ThermoFisher).

### RNA UREA PAGE and Northern blotting

Analysis of RNA by acrylamide gel electrophoresis in this study was performed using homemade 10 cm 8% (total RNA) or 16% (tRNA) TBE-urea gels (29:1 acrylamide:bis-acrylamide). Prior to electrophoresis, RNA was denatured by heating to 95°C in the presence of an equal volume of 2x RNA loading dye (95% v/v formamide, 0.025% bromophenol blue, 0.025% SDS, 10 mM EDTA pH 8.0). Gels were run using a mini-PROTEAN vertical electrophoresis system (BioRad) at 190 V for 35 minutes. Total RNA was visualized by staining with SYBR Gold diluted 1:10 000 in 10 mL 1x TBE for 10 minutes. Gels were imaged using a ChemiDoc instrument (BioRad).

For Northern blotting, RNA was transferred onto Zeta-probe quaternary amine nylon membranes (BioRad) in 1x TBE using the Mini Trans-Blot electrophoretic transfer system (BioRad) run at 0.4 A for 45 minutes. RNA was subsequently crosslinked to the membrane by exposure to UV for 1 minute using the UV lamp of a biological safety cabinet (10 cm distance between membrane and lamp). The membrane was then placed directly into 10 mL ULTRAhyb-oligo (ThermoFisher) hybridization buffer pre-heated to 42°C shaking at 60 RPM. After 10 minutes of pre-hybridization, biotinylated DNA probe was added to a final concentration of 10 nM and hybridization was allowed to proceed overnight. The membrane was subsequently washed twice for 30 minutes with 10 mL stringency wash buffer (2x SCC [ThermoFisher], 0.5% w/v SDS) at 42°C shaking at 60 RPM. The membrane was moved to room temperature, washed twice for 5 minutes each with 20 mL wash buffer (1x PBS pH 7.4, 0.5% w/v SDS), washed twice for 5 minutes each with 10 mL blocking buffer (2x SCC, 0.5% w/v SDS, 0.1% w/v I-block reagent [ThermoFisher]), and blocked for 30 minutes with 20 mL blocking buffer. Strepavidin-horseradish peroxidase was added (1:5000 v/v) and the membrane was incubated at room temperature with gentle shaking for 1 hour. The membrane was first washed with 20 mL blocking buffer for 15 minutes, then washed twice with 20 mL wash buffer. The blot was developed by the addition of Clarity Max ECL substrate (BioRad) and visualized using a ChemiDoc instrument (BioRad).

### S. aureus protein secretion assays

To collect proteins from *S. aureus* culture supernatant for Western blot analysis, overnight cultures of *S. aureus* strains harbouring the indicated pRAB11-derived vectors were diluted to OD600 of 0.1 in 50 mL TSB medium containing 10 μg/mL chloramphenicol and grown at 37°C shaking at 220 RPM. Anhydrotetracycline was added when cultures reached OD600 of 0.5, and cultures were allowed to continue growing until they reached OD600 8.0. Cells were pelleted by centrifugation at 6000 *g* for 20 minutes and trichloroacetic acid was added to the supernatant to a final concentration of 10% v/v. Supernatants were incubated rotating at 4°C overnight, and precipitated proteins were collected by centrifugation at 36 000 *g* for 30 minutes. Pellets were washed with 25 mL acetone, dried in a lamellar flow hood for 20 minutes, and resuspended in 200 μL 2.66x SDS-PAGE loading dye with 5.3 M urea. Samples were boiled for 10 minutes before analysis by Western blotting.

### Protein SDS-PAGE and Western blotting

Analysis of proteins by SDS-PAGE was performed using homemade 10 cm Tris-glycine SDS-PAGE gels containing 18% (w/v) acrylamide. Samples were boiled in the presence of 1x SDS PAGE loading dye at 95°C for 10 minutes, centrifuged at 21 000 *g* for 1 minute, and loaded onto a SDS-PAGE gel. Gels were run first for 12 minutes at 95 V, then for 40 minutes at 195 V. To visualize total proteins, gels were subsequently stained with Coomassie Brilliant Blue for 30 minutes at room temperature with gentle rotation and destained with 30% (v/v) methanol and 10% (v/v). For Western blotting, total protein was wet transferred from the SDS gel onto a nitrocellulose membrane (BioRad) using the Mini Trans-Blot electrophoresis system (BioRad). The transfer was run at 103 V for 33 minutes, after which the membrane was blocked in tris-buffered saline pH 7.2 containing 0.05% (v/v) Tween-20 (TBS-T) and 0.5% (w/v) non-fat milk (BioRad) for 30 minutes at room temperature with gentle shaking. For Western blot analysis of *S. aureus* supernatant, human serum was added (1:1000 dilution) to the blocking solution. Primary antibodies were added (1:5000) and the membrane was incubated for 1 hour at room temperature with gentle shaking. The membrane was washed 3x for 5 minutes each with 10 mL TBS-T and secondary antibody (anti-rabbit horseradish-peroxidase conjugate, MilliporeSigma) was added diluted 1:5000 in 10 mL TBS-T. The membrane was incubated for 45 minutes at room temperature with gentle shaking and washed 3x for 5 minutes each with 10 mL TBS-T. The blot was developed by the addition of Clarity Max ECL substrate (BioRad) and visualized using a ChemiDoc instrument (BioRad).

### Recombinant protein expression and purification

*E. coli* BL21 (DE3) pLysS (Novagen) strains harbouring expression vectors were grown in LB at 37°C shaking at 220 RPM overnight with appropriate antibiotics. 20 mL of starter culture was used to inoculate each 1 L LB, which was allowed to grow at 37°C shaking at 220 RPM to an OD600 of 0.6. IPTG was added at a final concentration of 1 mM to induce protein expression. The culture was subsequently incubated for 18 hours at 18°C shaking at 220 RPM before cells were harvested by centrifugation at 6000 *g* for 20 minutes. Cells were resuspended in lysis buffer (300 mM NaCl, 50 mM HEPES NaOH pH 7.5) and lysed by sonication. Lysates were clarified by centrifugation at 36 000 *g* for 45 minutes at 4°C and loaded onto a 1 mL gravity flow Ni-NTA agarose (Qiagen) column pre-equilibrated with 10 mL wash buffer (300 mM NaCl, 50 mM HEPES pH 7.5, 10 mM imidazole). The column was subsequently washed 3 times with 20 mL wash buffer before His6-tagged proteins were eluted by the addition of 4 mL elution buffer (300 mM NaCl, 50 mM HEPES pH 7.5, 400 mM imidazole). Eluted proteins were further purified by gel filtration on a HiLoad 16/600 superdex 200 size exclusion chromatography column (GE Healthcare) equilibrated with SEC buffer (20 mM HEPES pH 7.5, 150 mM NaCl). Protein purity was analyzed by SDS-PAGE stained with Coomassie Brilliant Blue R250. Purified proteins were sterilized by passing through a PES syringe filter with 0.22 μm pores (FroggaBio). Protein concentration was measured using a NanoDrop instrument (ThermoFisher), and proteins were concentrated to 50 μM using a 10 kDa molecular weight cutoff centrifugal filter device (MilliporeSigma). Proteins were used immediately or snap frozen in liquid nitrogen and stored at −80°C for later use.

### MIC determination

The MIC of purified AbpT*^Cc^*was determined by broth microdilution. 100 μL of the indicated 2-fold serial dilution of AbpT*^Cc^* alone, AbpT*^Cc^*and equimolar AbpP*^Cc^*, or an equivalent volume of buffer diluted in appropriate growth medium was loaded into a flat-bottom 96-well polystyrene plate. Overnight cultures of the indicated bacterial strains were diluted 10-fold, and 1 μL of this dilution was used to inoculate each 100 μL well (1:1000 subculture from original stationary phase culture). Plates were incubated in a stationary incubator at the appropriate growth temperature (supplementary table 1) for 18 hours before the OD600 was read using a a BioTek Synergy H1 multimode reader (Agilent). Assays were conducted in biological duplicate.

### Time-dependent bacterial killing assays

An overnight culture of the indicated strain was subcultured 1000-fold into 100 μL of appropriate medium containing purified AbpT*^Cc^*/AbpP*^Cc^*in equimolar amounts at 1.5x MIC or an equal volume of buffer (150 mM NaCl, 20 mM HEPES NaOH pH 7.5) in a flat-bottom 96-well polystyrene plate. At each time point, 10 μL was diluted in 90 μL growth medium, cells were pelleted by centrifugation at 4000xg for 15 minutes, resuspended in 100 μL growth medium and 10-fold serial dilutions were plated on appropriate growth medium plates for the indicated strain (supplementary table 1). Plates were incubated at the appropriate growth temperature and colonies were counted to calculate viable colony forming units (CFUs) per mL. Experiments were performed in biological triplicate.

To determine the effect of CCCP on AbpT*^Cc^* susceptibility, time-dependent killing assays were performed as above except that cultures were concomitantly treated with CCCP (5 μM for *Bacillus subtilis* and 100 μM for *Corynebacterium glutamicum*) or an equivalent volume of solvent (DMSO).

### ABP toxin proteolytic cleavage assays

The indicated AbpT and AbpP proteins were combined to a final concentration of 10 μM in a 12 μL reaction and incubated at 37°C for 30 minutes. Reactions were stopped by the addition of 4 μL 4x SDS-PAGE loading dye, samples were boiled at 95°C for 10 minutes and 6 μL of sample was run on an 18% acrylamide SDS-PAGE gel. Total proteins were visualized by staining with Coomassie Brilliant Blue R250.

For time-dependent protease cleavage assays, the indicated AbpT and AbpP proteins were combined in a 50 μL reaction and incubated at 37°C. At each of the indicated timepoints, a 6 μL aliquot of each reaction was quenched by the addition of 2 μL SDS-PAGE loading dye. Samples were boiled and 6 μL was run on an 18% acrylamide SDS-PAGE gel and visualized by staining with Coomassie Brilliant Blue R250.

### N-terminal sequencing

20 μM AbpT*^Cc^* was combined with an equimolar amount of AbpP*^Cc^* in a total reaction volume of 30 μL and incubated for 30 minutes at 37°C. 10 μL SDS-PAGE loading dye was added, the sample was boiled for 10 minutes and centrifuged for 1 minute at 21000xg to pellet protein aggregates. The sample was split across 2 wells of a Mini-PROTEAN pre-cast SDS-PAGE gel (BioRad), which was run at 200 V for 40 minutes. Total proteins were wet transferred to a PVDF membrane (BioRad) pre-soaked in 100% methanol using a Mini Trans-Blot electrophoretic transfer system (BioRad) run at 80V for 1 hour. The membrane was stained with Coomassie Brilliant Blue R250 and the band corresponding to the C-terminal fragment of AbpT*^Cc^* was excised. 5 cycles of N-terminal sequencing was performed by Protein Facility at Iowa State University. Sequencing results are provided in supplementary file 1.

### Bacterial strains and growth conditions

A complete list of strains and growth conditions used in this study can be found in supplementary table 1, and all strains are available upon request from the corresponding authors. *Escherichia coli* strain XL1 Blue (Novagen) and DC10b were used for plasmid maintenance and strain BL21 (DE3) pLysS (Novagen) was used for recombinant protein expression. *E. coli* strains were grown in lysogen broth (LB) medium (10 g/L tryptone, 5 g/L yeast extract, 10 g/L NaCl) at 37°C shaking at 220 revolutions per minute (RPM). *E. coli* cultures were supplemented with 50 μg/mL kanamycin, 100 μg/mL ampicillin, 25 μg/mL chloramphenicol, 200 μg/mL trimethoprim, 15 μg/mL gentamicin, 0.1% L-rhamnose, 0.1% L-arabinose, and 1 mM isopropyl β-D-1-thiogalactopyranoside, as appropriate. *Staphylococcus aureus* cultures were grown in tryptic soy broth (TSB) medium (Bacto) at 37°C shaking at 220 revolutions per minute (RPM). *S. aureus* cultures were supplemented with 10 μg/mL chloramphenicol and 500 ng/mL anhydrotetracycline as appropriate.

### DNA manipulation and plasmid construction

All expression vectors were constructed using standard restriction enzyme-based cloning procedures (49). Primers were synthesized by Integrated DNA Technologies. Phusion polymerase, restriction enzymes, and T4 DNA ligase were obtained from New England Biolabs. Whole plasmid sequencing was performed by Plasmidsaurus using Oxford Nanopore Technology with custom analysis and annotation. Oligonucleotide sequences and plasmids used in this study are provided in supplementary tables 2 and 3, respectively.

### Preparation of concentrated supernatant for bacterial growth inhibition assays

*S. aureus* strains derived from the sequenced strain USA300 harbouring the indicated pRAB11 vectors were grown overnight at 37°C shaking at 220 RPM in 5 mL TSB supplemented with 10 μg/mL chloramphenicol. Overnight cultures were diluted to OD600 of 0.1 in 500 mL TSB with 10 μg/mL chloramphenicol and grown at 37°C shaking at 220 RPM to an OD600 of 0.5. Anhydrotetracycline was added to a final concentration of 500 ng/mL to induce gene expression. Upon reaching OD600 of 8.0, cultures were pelleted at 6000 *g* for 20 minutes and protein was precipitated from culture supernatant by the addition of ammonium sulfate to a final concentration of 80% (w/v). Supernatant was stirred overnight at 4°C and subsequently centrifuged at 36 000 *g* for 45 minutes. Precipitated supernatant was resuspended in 10 mL 1x phosphate buffered saline (PBS) pH 7.4 and dialyzed twice against 2 L 1x PBS at room temperature for 1 hour each. Dialyzed protein was further concentrated using a 10 kDa molecular weight cutoff centrifugal filter device (MilliporeSigma) to a final volume of 2 mL (250-fold concentration of original culture supernatant). Concentrated protein was snap frozen in liquid nitrogen and stored at −80°C before use.

### Bacterial growth inhibition screening

Strains used in this assay, their sources, and growth conditions are listed in supplementary table 1. Starting cultures were diluted to an OD of 0.05 in 90 μL of appropriate media in adjacent wells of a 96 well plate. 10 μL of concentrated culture supernatant (see above) collected from *S. aureus* cultures expressing either the indicated ABP system or an empty vector control was added to each well. Plates were subsequently incubated in growth conditions appropriate for each strain for 8 hours, and OD600 was measured using a BioTek Synergy H1 multimode reader (Agilent). Two biological replicates of the screen were conducted, and the p-values were calculated using an unpaired T-test comparing mean OD600 of the culture treated with ABP supernatant compared with the culture treated with control supernatant.

### Bacterial growth curves

To validate initial hits from the ABP susceptibility screen, stationary phase cultures of the indicated bacterial strains grown in appropriate media were subcultured 1:100 into 96 well plates containing 90 μL of appropriate media, to which 10 μL concentrated ABP or control culture supernatant was added. Cultures were grown at the indicated temperatures in the presence or absence of shaking in a BioTek Synergy H1 multimode reader (Agilent) and the absorbance at 600 nm was measured every 30 minutes.

To determine the growth inhibition activity of purified AbpT proteins, experiments were conducted as above, but the indicated purified proteins were added at 1.5x their MIC instead of concentrated culture supernatant.

### E. coli toxicity experiments

Cultures of *E. coli* XL1 Blue harbouring the indicated plasmids were grown overnight in 2 mL LB supplemented with appropriate antibiotics at 37°C shaking at 220 RPM. Overnight cultures were normalized to an OD600 of 1.0 and were diluted in 10-fold series in a 96 well microtiter plate. 6 μL of 10^-1^-10^-6^ dilutions were spotted on LB 1.5% (w/v) agar containing 0.1% (w/v) L-rhamnose or 0.1% (w/v) L-arabinose to induce toxin expression from pSCrhaB2-CV- or pBAD33-derived vectors, respectively, or 1 mM IPTG to induce immunity expression from pPSV39-derived vectors. Plates were incubated at 37°C for 18 hours and photographed.

### RNA extraction

Cultures of *E. coli* XL1 Blue harbouring the indicated plasmids were grown overnight in 2 mL LB supplemented with appropriate antibiotics at 37°C shaking at 220 RPM. Cultures were diluted to OD600 0.1 in 25 mL LB supplemented with appropriate antibiotics and grown at 37°C shaking at 220 RPM for approximately 2 hours to OD600 of 0.5. Toxin expression was induced by the addition of L-rhamnose to a final concentration of 0.1% (w/v). After the indicated timepoints had elapsed, 5 mL culture was collected by centrifugation at 4000 *g* for 10 minutes at room temperature and the pellet was resuspended in 1 mL TRIzol reagent (ThermoFisher). Total RNA was extracted using a standard protocol (50). Briefly, *E. coli* resuspended in TRIzol reagent was vortexed for 30 seconds and incubated at room temperature for 5 minutes before the addition of 200 μL chloroform. The mixture was vortexed for 30 seconds and incubated on ice for 2 minutes before centrifugation at 14 000 *g* for 15 minutes at 4°C. 500 μL of the upper aqueous phase was pipetted into 500 μL of ice cold isopropanol, vortexed for 30 seconds, incubated on ice for 20 minutes, and centrifuged at 21 000 *g* for 15 minutes at 4°C. The resulting RNA pellet was washed three times with 1 mL ice cold 75% (v/v) ethanol and dried under a vacuum. The RNA pellet was resuspended in 100 μL 1x DNAse I buffer (NEB) to which 4 units RNAse-free DNAse I (NEB) was added. The reaction was incubated for 30 minutes at 37°C and RNA was subsequently extracted using a Monarch Spin RNA cleanup kit (NEB) following the manufacturer’s instructions. RNA was quantified using a NanoDrop spectrophotometer (ThermoFisher) and 500 ng of total RNA was analyzed by UREA PAGE stained with SYBR Gold (ThermoFisher).

The method above was used to extract RNA from *Bacillus subtilis* treated with purified AbpT, with the following modifications. An isolated colony of *B. subtilis* strain PY79 was collected from a LB plate containing 1.5% (w/v) agar and grown to OD600 of 0.5 in 10 mL LB at 37°C shaking at 220 RPM. The 10 mL culture was subsequently split into 2x 5 mL cultures; one culture was treated with purified AbpT*^Cc^* and AbpP*^Cc^* to a final concentration of 1 μM each and the other 5 mL culture was treated with an equal volume of sterile buffer (150 mM NaCl, 20 mM HEPES NaOH pH 7.5). Cultures were subsequently incubated at 37°C shaking at 220 RPM for 1 h before cells were collected by centrifugation at 4000 *g* at room temperature for 10 minutes. Pellets were washed 3x with 1 mL 1x PBS pH 7.4. Pellets were resuspended in 200 μL 1x PBS pH 7.4 containing 50 mg/mL hen egg lysozyme and incubated for 20 minutes at 37°C. 1 mL TRIzol reagent was subsequently added, and RNA extraction was completed as described above.

### RNAse activity assays

To determine the ribonuclease activity of AbpT*^Cc^* and AbpT*^Pm^ in vitro*, 1 μM of the indicated purified proteins was incubated with 2000 ng of total *E. coli* RNA extracted from *E. coli* strain MG1655 using the method described above or 2000 ng purified *E. coli* tRNA (Roche) in a 6 μL reaction volume in buffer containing 150 mM NaCl, 20 mM HEPES NaOH pH 7.5. Reactions were allowed to proceed for 20 minutes at 37°C and were quenched by the addition of an equal volume of 2x RNA loading dye (95% v/v formamide, 0.025% bromophenol blue, 0.025% SDS, 10 mM EDTA pH 8.0). Samples were subsequently heated to 95°C for 10 minutes and rapidly cooled by submerging in ice water. 500 ng total RNA was analyzed by UREA PAGE stained with SYBR Gold (ThermoFisher).

### RNA UREA PAGE and Northern blotting

Analysis of RNA by acrylamide gel electrophoresis in this study was performed using homemade 10 cm 8% (total RNA) or 16% (tRNA) TBE-urea gels (29:1 acrylamide:bis-acrylamide). Prior to electrophoresis, RNA was denatured by heating to 95°C in the presence of an equal volume of 2x RNA loading dye (95% v/v formamide, 0.025% bromophenol blue, 0.025% SDS, 10 mM EDTA pH 8.0). Gels were run using a mini-PROTEAN vertical electrophoresis system (BioRad) at 190 V for 35 minutes. Total RNA was visualized by staining with SYBR Gold diluted 1:10 000 in 10 mL 1x TBE for 10 minutes. Gels were imaged using a ChemiDoc instrument (BioRad).

For Northern blotting, RNA was transferred onto Zeta-probe quaternary amine nylon membranes (BioRad) in 1x TBE using the Mini Trans-Blot electrophoretic transfer system (BioRad) run at 0.4 A for 45 minutes. RNA was subsequently crosslinked to the membrane by exposure to UV for 1 minute using the UV lamp of a biological safety cabinet (10 cm distance between membrane and lamp). The membrane was then placed directly into 10 mL ULTRAhyb-oligo (ThermoFisher) hybridization buffer pre-heated to 42°C shaking at 60 RPM. After 10 minutes of pre-hybridization, biotinylated DNA probe was added to a final concentration of 10 nM and hybridization was allowed to proceed overnight. The membrane was subsequently washed twice for 30 minutes with 10 mL stringency wash buffer (2x SCC [ThermoFisher], 0.5% w/v SDS) at 42°C shaking at 60 RPM. The membrane was moved to room temperature, washed twice for 5 minutes each with 20 mL wash buffer (1x PBS pH 7.4, 0.5% w/v SDS), washed twice for 5 minutes each with 10 mL blocking buffer (2x SCC, 0.5% w/v SDS, 0.1% w/v I-block reagent [ThermoFisher]), and blocked for 30 minutes with 20 mL blocking buffer. Strepavidin-horseradish peroxidase was added (1:5000 v/v) and the membrane was incubated at room temperature with gentle shaking for 1 hour. The membrane was first washed with 20 mL blocking buffer for 15 minutes, then washed twice with 20 mL wash buffer. The blot was developed by the addition of Clarity Max ECL substrate (BioRad) and visualized using a ChemiDoc instrument (BioRad).

### S. aureus protein secretion assays

To collect proteins from *S. aureus* culture supernatant for Western blot analysis, overnight cultures of *S. aureus* strains harbouring the indicated pRAB11-derived vectors were diluted to OD600 of 0.1 in 50 mL TSB medium containing 10 μg/mL chloramphenicol and grown at 37°C shaking at 220 RPM. Anhydrotetracycline was added when cultures reached OD600 of 0.5, and cultures were allowed to continue growing until they reached OD600 8.0. Cells were pelleted by centrifugation at 6000 *g* for 20 minutes and trichloroacetic acid was added to the supernatant to a final concentration of 10% v/v. Supernatants were incubated rotating at 4°C overnight, and precipitated proteins were collected by centrifugation at 36 000 *g* for 30 minutes. Pellets were washed with 25 mL acetone, dried in a lamellar flow hood for 20 minutes, and resuspended in 200 μL 2.66x SDS-PAGE loading dye with 5.3 M urea. Samples were boiled for 10 minutes before analysis by Western blotting.

### Protein SDS-PAGE and Western blotting

Analysis of proteins by SDS-PAGE was performed using homemade 10 cm Tris-glycine SDS-PAGE gels containing 18% (w/v) acrylamide. Samples were boiled in the presence of 1x SDS PAGE loading dye at 95°C for 10 minutes, centrifuged at 21 000 *g* for 1 minute, and loaded onto a SDS-PAGE gel. Gels were run first for 12 minutes at 95 V, then for 40 minutes at 195 V. To visualize total proteins, gels were subsequently stained with Coomassie Brilliant Blue for 30 minutes at room temperature with gentle rotation and destained with 30% (v/v) methanol and 10% (v/v). For Western blotting, total protein was wet transferred from the SDS gel onto a nitrocellulose membrane (BioRad) using the Mini Trans-Blot electrophoresis system (BioRad). The transfer was run at 103 V for 33 minutes, after which the membrane was blocked in tris-buffered saline pH 7.2 containing 0.05% (v/v) Tween-20 (TBS-T) and 0.5% (w/v) non-fat milk (BioRad) for 30 minutes at room temperature with gentle shaking. For Western blot analysis of *S. aureus* supernatant, human serum was added (1:1000 dilution) to the blocking solution. Primary antibodies were added (1:5000) and the membrane was incubated for 1 hour at room temperature with gentle shaking. The membrane was washed 3x for 5 minutes each with 10 mL TBS-T and secondary antibody (anti-rabbit horseradish-peroxidase conjugate, MilliporeSigma) was added diluted 1:5000 in 10 mL TBS-T. The membrane was incubated for 45 minutes at room temperature with gentle shaking and washed 3x for 5 minutes each with 10 mL TBS-T. The blot was developed by the addition of Clarity Max ECL substrate (BioRad) and visualized using a ChemiDoc instrument (BioRad).

### Recombinant protein expression and purification

*E. coli* BL21 (DE3) pLysS (Novagen) strains harbouring expression vectors were grown in LB at 37°C shaking at 220 RPM overnight with appropriate antibiotics. 20 mL of starter culture was used to inoculate each 1 L LB, which was allowed to grow at 37°C shaking at 220 RPM to an OD600 of 0.6. IPTG was added at a final concentration of 1 mM to induce protein expression. The culture was subsequently incubated for 18 hours at 18°C shaking at 220 RPM before cells were harvested by centrifugation at 6000 *g* for 20 minutes. Cells were resuspended in lysis buffer (300 mM NaCl, 50 mM HEPES NaOH pH 7.5) and lysed by sonication. Lysates were clarified by centrifugation at 36 000 *g* for 45 minutes at 4°C and loaded onto a 1 mL gravity flow Ni-NTA agarose (Qiagen) column pre-equilibrated with 10 mL wash buffer (300 mM NaCl, 50 mM HEPES pH 7.5, 10 mM imidazole). The column was subsequently washed 3 times with 20 mL wash buffer before His6-tagged proteins were eluted by the addition of 4 mL elution buffer (300 mM NaCl, 50 mM HEPES pH 7.5, 400 mM imidazole). Eluted proteins were further purified by gel filtration on a HiLoad 16/600 superdex 200 size exclusion chromatography column (GE Healthcare) equilibrated with SEC buffer (20 mM HEPES pH 7.5, 150 mM NaCl). Protein purity was analyzed by SDS-PAGE stained with Coomassie Brilliant Blue R250. Purified proteins were sterilized by passing through a PES syringe filter with 0.22 μm pores (FroggaBio). Protein concentration was measured using a NanoDrop instrument (ThermoFisher), and proteins were concentrated to 50 μM using a 10 kDa molecular weight cutoff centrifugal filter device (MilliporeSigma). Proteins were used immediately or snap frozen in liquid nitrogen and stored at −80°C for later use.

### MIC determination

The MIC of purified AbpT*^Cc^*was determined by broth microdilution. 100 μL of the indicated 2-fold serial dilution of AbpT*^Cc^* alone, AbpT*^Cc^*and equimolar AbpP*^Cc^*, or an equivalent volume of buffer diluted in appropriate growth medium was loaded into a flat-bottom 96-well polystyrene plate. Overnight cultures of the indicated bacterial strains were diluted 10-fold, and 1 μL of this dilution was used to inoculate each 100 μL well (1:1000 subculture from original stationary phase culture). Plates were incubated in a stationary incubator at the appropriate growth temperature (supplementary table 1) for 18 hours before the OD600 was read using a a BioTek Synergy H1 multimode reader (Agilent). Assays were conducted in biological duplicate.

### Time-dependent bacterial killing assays

An overnight culture of the indicated strain was subcultured 1000-fold into 100 μL of appropriate medium containing purified AbpT*^Cc^*/AbpP*^Cc^*in equimolar amounts at 1.5x MIC or an equal volume of buffer (150 mM NaCl, 20 mM HEPES NaOH pH 7.5) in a flat-bottom 96-well polystyrene plate. At each time point, 10 μL was diluted in 90 μL growth medium, cells were pelleted by centrifugation at 4000xg for 15 minutes, resuspended in 100 μL growth medium and 10-fold serial dilutions were plated on appropriate growth medium plates for the indicated strain (supplementary table 1). Plates were incubated at the appropriate growth temperature and colonies were counted to calculate viable colony forming units (CFUs) per mL. Experiments were performed in biological triplicate.

To determine the effect of CCCP on AbpT*^Cc^* susceptibility, time-dependent killing assays were performed as above except that cultures were concomitantly treated with CCCP (5 μM for *Bacillus subtilis* and 100 μM for *Corynebacterium glutamicum*) or an equivalent volume of solvent (DMSO).

### ABP toxin proteolytic cleavage assays

The indicated AbpT and AbpP proteins were combined to a final concentration of 10 μM in a 12 μL reaction and incubated at 37°C for 30 minutes. Reactions were stopped by the addition of 4 μL 4x SDS-PAGE loading dye, samples were boiled at 95°C for 10 minutes and 6 μL of sample was run on an 18% acrylamide SDS-PAGE gel. Total proteins were visualized by staining with Coomassie Brilliant Blue R250.

For time-dependent protease cleavage assays, the indicated AbpT and AbpP proteins were combined in a 50 μL reaction and incubated at 37°C. At each of the indicated timepoints, a 6 μL aliquot of each reaction was quenched by the addition of 2 μL SDS-PAGE loading dye. Samples were boiled and 6 μL was run on an 18% acrylamide SDS-PAGE gel and visualized by staining with Coomassie Brilliant Blue R250.

### N-terminal sequencing

20 μM AbpT*^Cc^* was combined with an equimolar amount of AbpP*^Cc^* in a total reaction volume of 30 μL and incubated for 30 minutes at 37°C. 10 μL SDS-PAGE loading dye was added, the sample was boiled for 10 minutes and centrifuged for 1 minute at 21000xg to pellet protein aggregates. The sample was split across 2 wells of a Mini-PROTEAN pre-cast SDS-PAGE gel (BioRad), which was run at 200 V for 40 minutes. Total proteins were wet transferred to a PVDF membrane (BioRad) pre-soaked in 100% methanol using a Mini Trans-Blot electrophoretic transfer system (BioRad) run at 80V for 1 hour. The membrane was stained with Coomassie Brilliant Blue R250 and the band corresponding to the C-terminal fragment of AbpT*^Cc^* was excised. 5 cycles of N-terminal sequencing were performed by the Protein Facility at Iowa State University. Sequencing results are provided in supplementary file 1.

## Acknowledgements

The authors thank Brian Coombes, Mark McBride, Michael Surette, Joshua Woodward, and Gerard Wright for gifts of bacterial strains; Polyniki Gkragkopoulou and Shehryar Ahmad for critical feedback on the manuscript; and members of the Whitney Lab for helpful discussions. J.C. is supported by a Canada Graduate Scholarship from the Canadian Institutes of Health Research (CIHR). S.R.G. is supported by a postdoctoral fellowship from the European Molecular Biology Organization (EMBO). JCW is the Canada Research Chair in Molecular Microbiology and holds an Investigators in the Pathogenesis of Infectious Disease Award from the Burroughs Wellcome Fund. This work was supported by a Discovery Grant from the Natural Sciences and Engineering Research Council of Canada (NSERC) (RGPIN-2025-06280 to JCW).

## Author contributions

J.C. designed experiments, performed all experiments, analyzed data, prepared figures, and wrote the manuscript. S.R.G. edited the manuscript and conceptualized the study. J.C.W. conceptualized the study, secured funding, and edited the manuscript.

## Competing interests

The authors declare no competing interests.

## Materials & Correspondence

All materials used in this study are available from the corresponding author John C. Whitney.

## Notes

### Competing Interest Statement

The authors have declared no competing interest.

