## Supplementary Material for "Proteolytically activated antibacterial toxins inhibit the growth of diverse Gram-positive bacteria"

Figure S1-S10
Table S1: Bacterial strains used for ABP susceptibility screening
Table S2: Strains used in this study
Table S3: Plasmids used in this study
Supplementary References

Keywords:
*To whom correspondence should be addressed: John C Whitney or Stephen R Garrett
Telephone – (+1) 905-525-9140

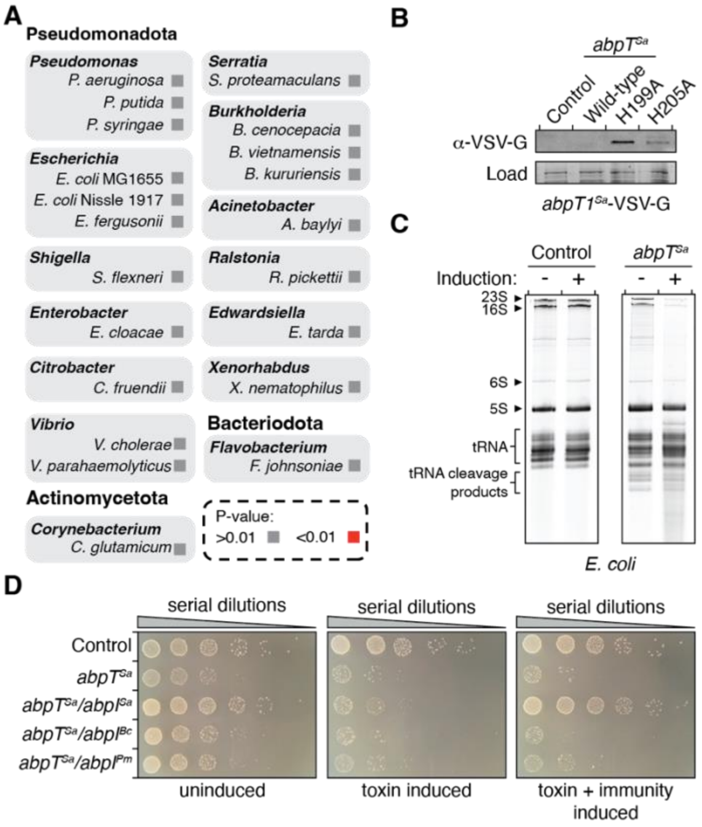


**Figure S1: Characterization of AbpT$^{Sa}$**. A) Subset of ABP susceptibility screening results. P values calculated from a one-tailed homoscedastic T test comparing growth of the indicated bacterial strain grown in the presence or absence of supernatant from an ABP-expressing *S. aureus* strain. B) Western blot analysis of *E. coli* expressing the indicated *abpT*$^{Sa}$ variants. An abundant protein species visualized on SDS PAGE stained with Coomassie Brilliant Blue R250 is used as a loading control. C) Urea PAGE analysis of total RNA extracted from *E. coli* harbouring a plasmid encoding *abpT*$^{Sa}$ or an empty vector control in the presence or absence of inducer, as indicated. Total RNA was visualized by staining with SYBR Gold. D) Growth of *E. coli* harbouring plasmids encoding the indicated genes under the control of distinct promoters on solid media containing separate inducers for each gene.

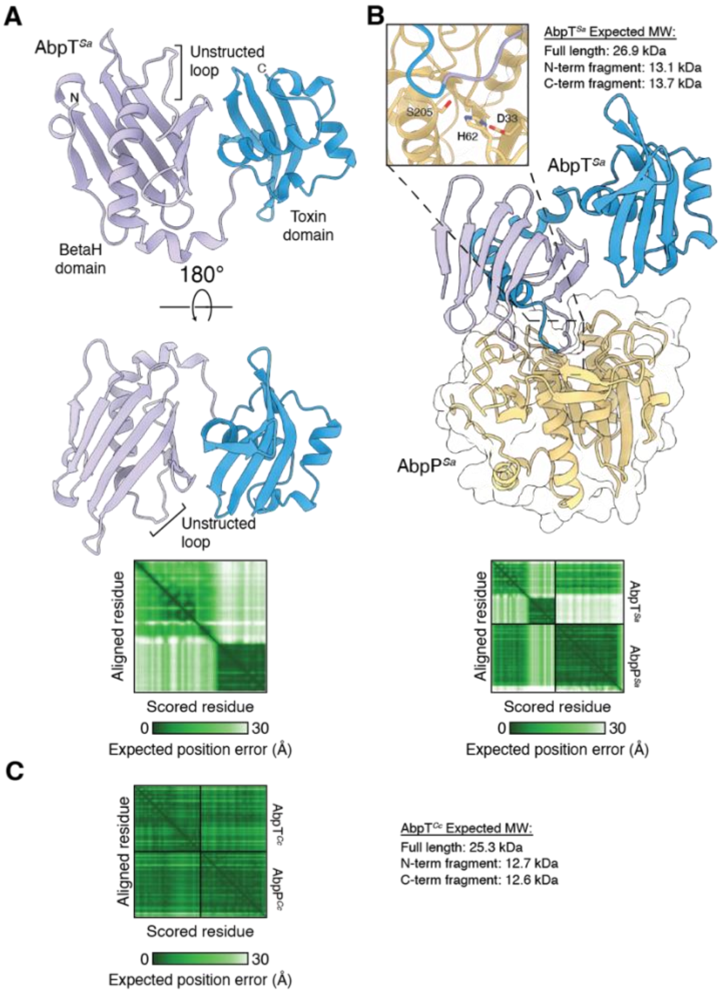


**Figure S2: AlphaFold3 modelling of AbpT alone and in complex with AbpP.** A) AlphaFold3 model of AbpT$^{Sa}$. The N-terminal BetaH domain and C-terminal toxin domain are indicated. Predicted aligned error plot is provided below. B) AlphaFold3 model of the complex formed between AbpT$^{Sa}$ and its associated protease AbpP$^{Sa}$. Predicted aligned error plot is provided below. C) AlphaFold3 predicted aligned error plot of the complex formed between AbpT$^{Cc}$ and AbpP$^{Cc}$.

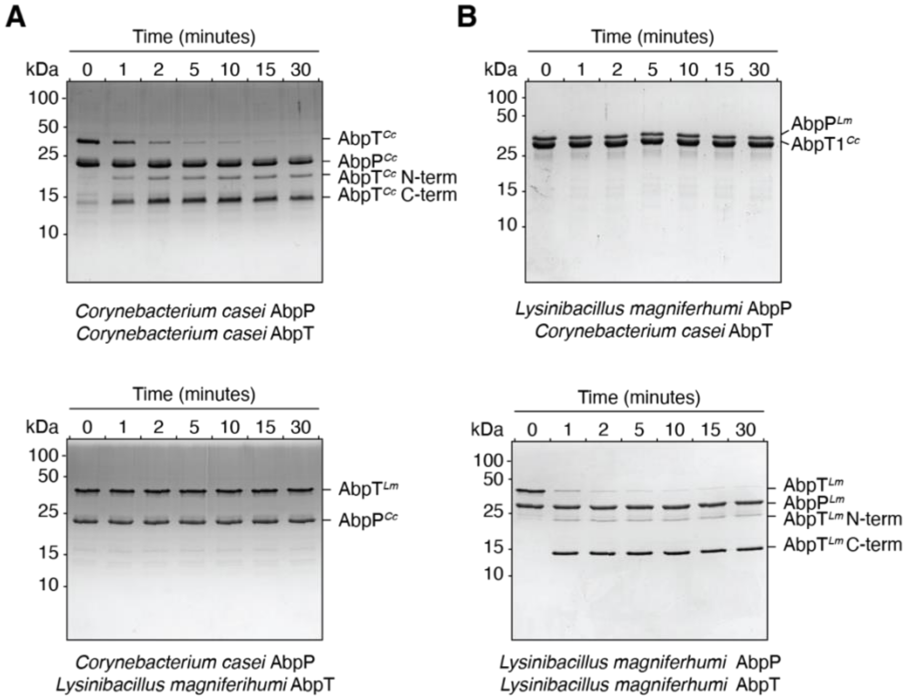


**Figure S3: AbpP proteases are specific for their cognate AbpT proteins.** A) SDS-PAGE analysis of AbpP$^{Cc}$ incubated with AbpT$^{Cc}$ or AbpT$^{Lm}$ for the indicated lengths of time. B) SDS-PAGE analysis of the AbpP$^{Lm}$ incubated with AbpT$^{Cc}$ or AbpT$^{Lm}$ for the indicated lengths of time.

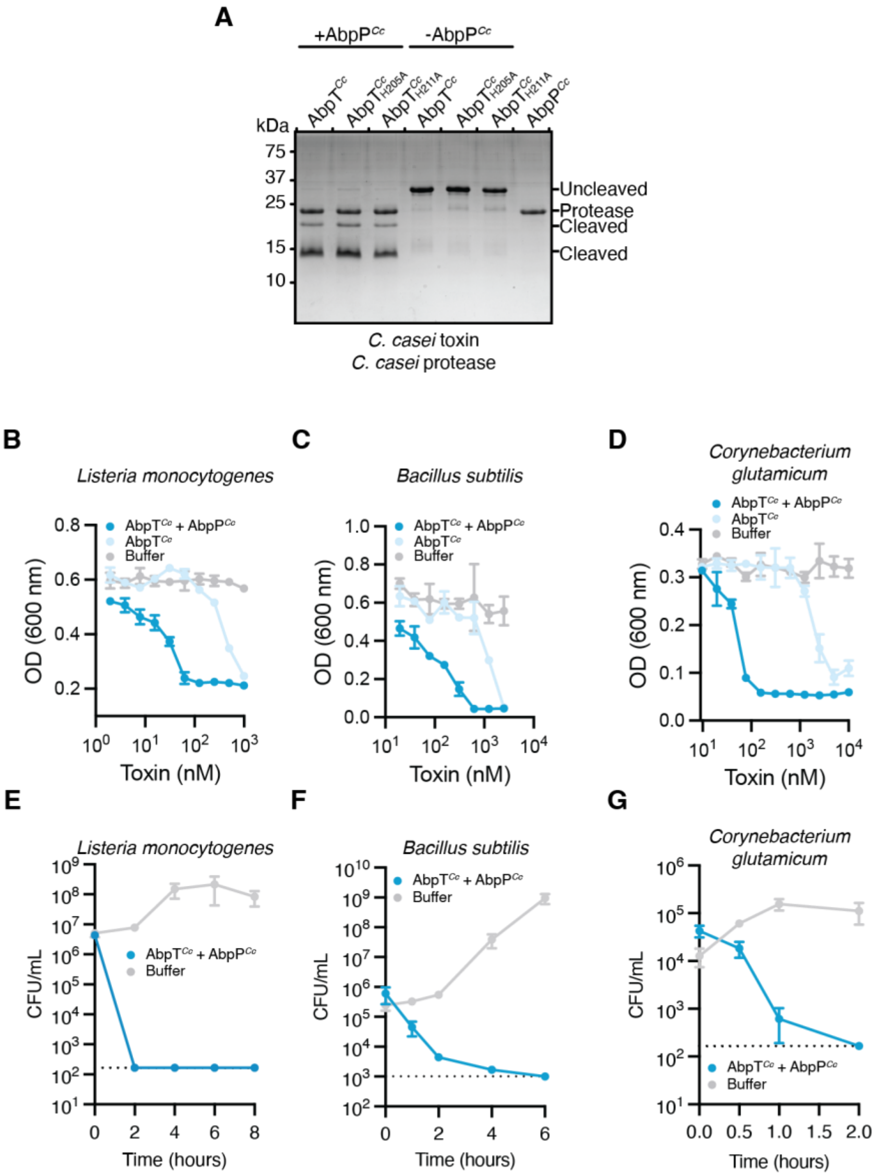


**Figure S4: Purified AbpT proteins display potent bactericidal activity against diverse Gram-positive bacteria.** A) SDS-PAGE analysis of the indicated AbpT$^{Cc}$ variants following cleavage by AbpT$^{Cc}$. B-D) MIC assays of AbpT$^{Cc}$ in the presence or absence of equimolar AbpP$^{Cc}$ against the indicated bacterial species. Data are presented as mean+/-SEM, n=2 biological replicates. E-G) Surviving colony forming units following incubation with proteolytically cleaved AbpT$^{Cc}$ at a concentration 1.5x MIC. The dashed line denotes the limit of detection. Data are presented as mean+SEM, n=3 biological replicates.

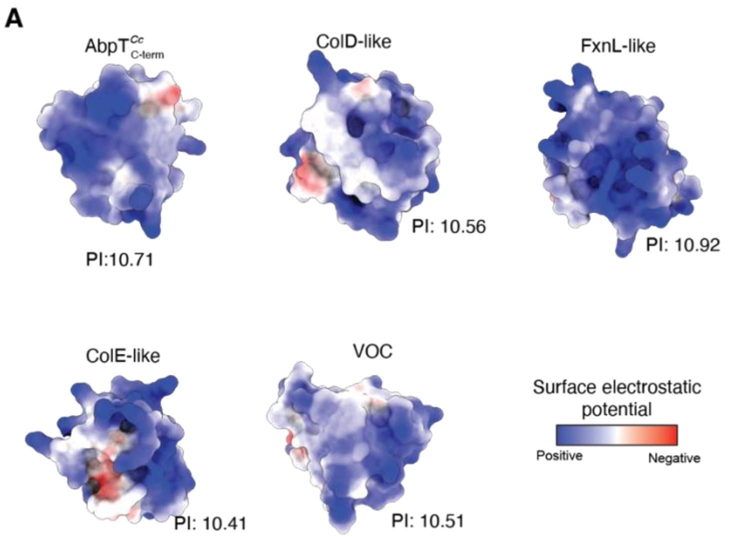


**Figure S5: AbpT toxin domains are highly cationic despite their structural and functional diversity.** A) Electrostatic surface representation of the representative ABP C-terminal toxin domains modelled using AlphaFold3. Predicted toxin activities and theoretical isoelectric points are indicated. These proteins are predicted to adopt the Colicin D, (ColD-like) Colicin E (ColE-like), vicinal oxygen chelate (VOC), or frataxin (FxnL-like) folds, as indicated.

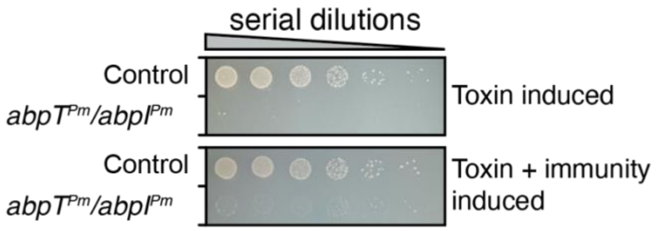


**Figure S6: AbpI$^{Pm}$ protects *E. coli* from growth inhibition by AbpT$^{Pm}$.** A) Growth of *E. coli* harbouring plasmids encoding the indicated genes under the control of distinct promoters on solid media containing separate inducers for each gene.

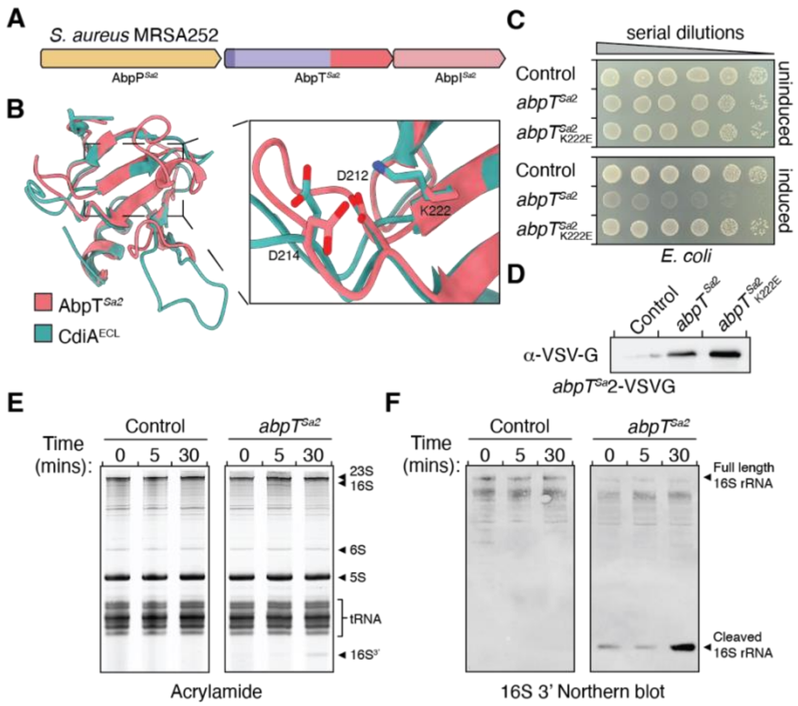


**Figure S7: An AbpT toxin encoded by *S. aureus* cleaves 16S ribosomal RNA**. A) Schematic representative of the ABP system encoded by the epidemic methicillin-resistant *S. aureus* strain MRSA252. B) AlphaFold3 model of the C-terminal BECR domain of AbpT$^{Sa2}$ overlaid with the C-terminal domain of CdiA$^{ECL}$ (PDB 4NTQ), an antibacterial toxin secreted by *Enterobacter cloacae* that targets the 16S ribosomal RNA. C) Viability of *E. coli* harbouring plasmids encoding the indicated *abpT*$^{Sa2}$ variants grown on solid media in the presence or absence of inducer. D) Western blot analysis of *E. coli* expressing the indicated *abpT*$^{Sa2}$ variants. E) Urea PAGE analysis of total RNA extracted from *E. coli* harbouring a plasmid encoding *abpT*$^{Sa}$ or an empty vector control following induction for the indicated lengths of time. Total RNA was visualized by staining with SYBR Gold. F) Northern blot analysis of the RNA in E) using a probe specific for the 3' end of the 16S ribosomal RNA.

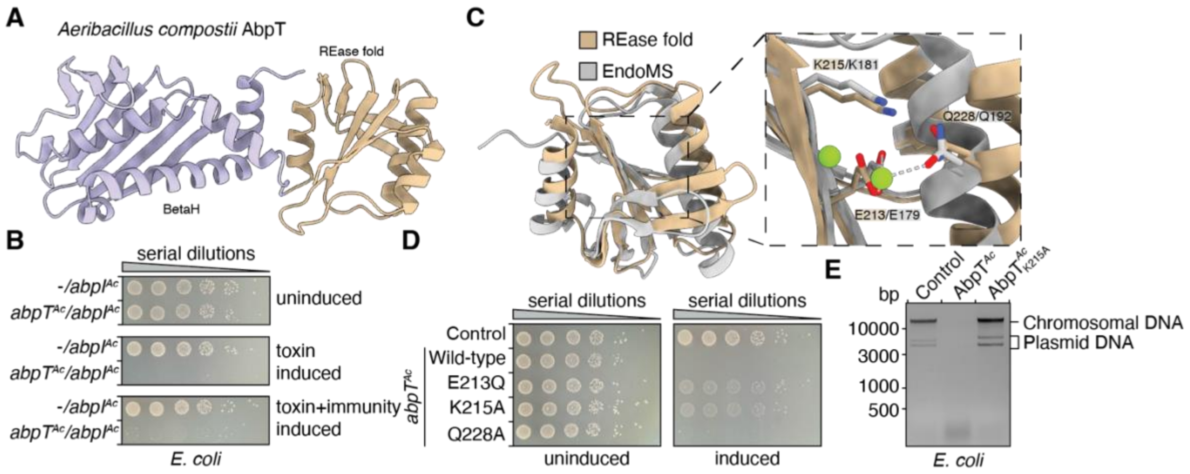


**Figure S8: An AbpT toxin encoded by *Aeribacillus compostii* is a DNAse of the restriction endonuclease superfamily** A) AlphaFold3 predicted structure of *Aeribacillus compostii* AbpT (AbpT$^{Ac}$). The N-terminal BetaH domain is highlighted in purple, and the C-terminal restriction endonuclease (REase) domain is highlighted in red. B) Growth of *E. coli* harbouring plasmids encoding *abpT$^{Ac}$* and *abpI$^{Ac}$* under the control of distinct promoters on solid media containing separate inducers for each gene, as indicated. C) AlphaFold3 model of the C-terminal REase domain of AbpT$^{Ac}$ overlaid with the restriction endonuclease EndoMS (PDB 5GKE). Residues forming the conserved magnesium-binding active site are highlighted in the insert. D) Viability of *E. coli* harbouring plasmids encoding the indicated *abpT$^{Ac}$* variants grown on solid media in the presence or absence of inducer. E) Genomic DNA extracted from *E. coli* cultures expressing the indicated *abpT$^{Ac}$* variants or an empty vector control.

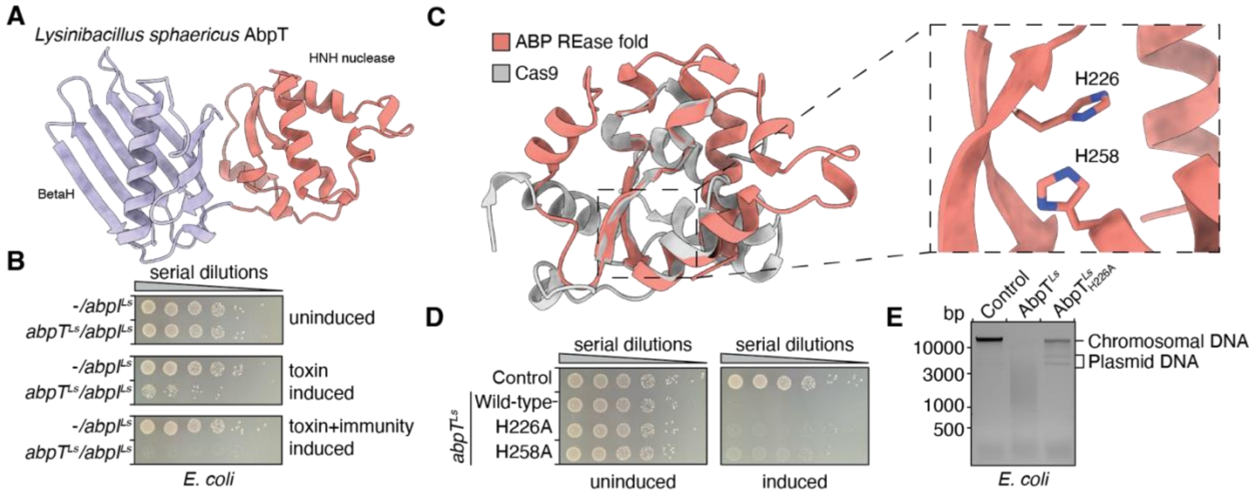


**Figure S9: An AbpT toxin encoded by *Lysinibacillus sphaericus* acts as a DNAse of the HNH superfamily** A) AlphaFold3 predicted structure of *Lysinibacillus sphaericus* AbpT (AbpT$^{Ls}$). The N-terminal BetaH domain is highlighted in purple, and the C-terminal toxin domain is highlighted in red. B) Growth of *E. coli* harbouring plasmids encoding *abpT*$^{Ls}$ and *abpI*$^{Ls}$ under the control of distinct promoters on solid media containing separate inducers for each gene, as indicated. C) AlphaFold3 model of the C-terminal HNH domain of AbpT$^{Ls}$ overlaid with the HNH nuclease domain of a CRISPR-associated nuclease (Cas) 9 enzyme encoded by *Geobacillus stearothermophilus* (PDB 8F43). A conserved pair of histidine residues in AbpT$^{Ls}$ is highlighted in the insert. D) Viability of *E. coli* harbouring plasmids encoding the indicated *abpT*$^{Ls}$ variants grown on solid media in the presence or absence of inducer. E) Genomic DNA extracted from *E. coli* cultures expressing the indicated *abpT*$^{Ls}$ variants or an empty vector control.

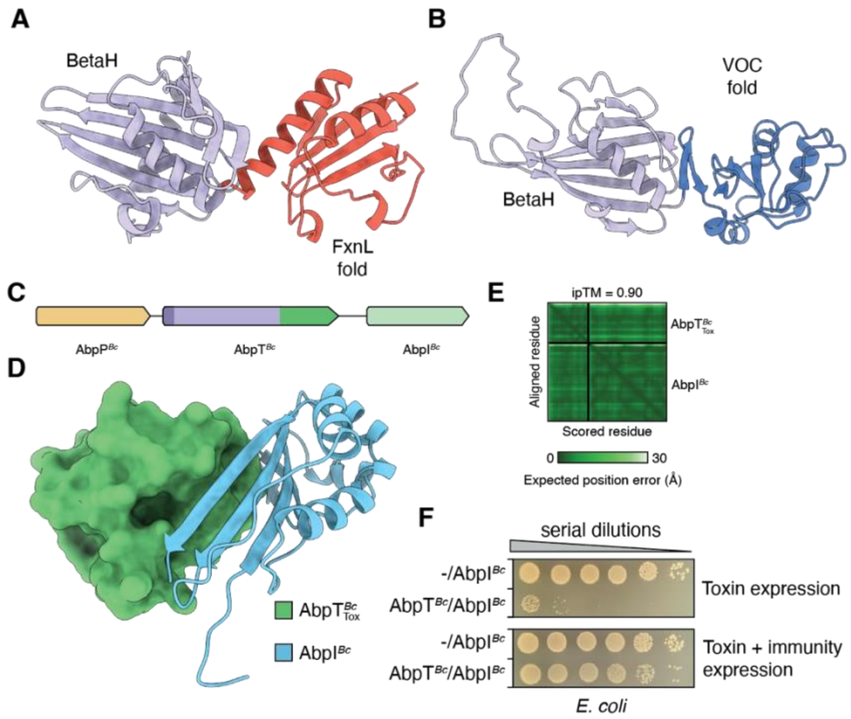


**Figure S10: ABP systems encode structurally unique toxins** A-B) AlphaFold3 models of representative AbpT proteins containing C-terminal toxins domains of unknown function predicted to adopt the frataxin-like (FxnL) or vicinal oxygen chelate (VOC) fold. The BetaH domains are coloured in purple. C) Schematic representative of the ABP system encoded by the soil bacterium *Bacillus cereus* strain ATCC 14579. D) AlphaFold3 prediction of the complex formed between the C-terminal domain of AbpT$^{Bc}$ and AbpI$^{Bc}$. The C-terminal AbpT$^{Bc}$ domain is predicted to adopt the uncharacterized Ntox33 domain fold and therefore likely acts by a unique mechanism. E) AlphaFold3 predicted aligned error plot for the AbpT$^{Bc}$/AbpI$^{Bc}$ complex shown in (D). F) Growth of *E. coli* harbouring plasmids encoding the indicated genes under the control of distinct promoters on solid media containing separate inducers for each gene.

**Table S1:** Bacterial strains used for ABP susceptibility screening

| Phylum | Genus and species | Strain | Medium | Temperat ure | Source |
|---|---|---|---|---|---|
| Actinomycetota | *Corynebacterium glutamicum* | MB001 | BHI | 30 | (1) |
| Bacillota | *Bacillus cereus* | ATCC 14579 | LB | 37 | (2) |
| Bacillota | *Bacillus subtilis* | JH642 | LB | 37 | (3) |
| Bacillota | *Bacillus subtilis* | PY79 | LB | 37 | (4) |
| Bacillota | *Bacillus subtilis* | 168 | LB | 37 | (5) |
| Bacillota | *Enterococcus faecalis* | OG1RF | THB | 37 | (6) |
| Bacillota | *Enterococcus faecalis* | V583 | THB | 37 | (7) |
| Bacillota | *Enterococcus faecalis* | ATCC 4200 | THB | 37 | (8) |
| Bacillota | *Enterococcus faecalis* | T1 | THB | 37 | (9) |
| Bacillota | *Enterococcus moraviensis* | BAA 383 | THB | 37 | (10) |
| Bacillota | *Listeria innocua* | | BHI | 37 | Gift from Joshua Woodward |
| Bacillota | *Lactococcus lactis* | GC982 | THY | 37 | Gift from Michael Surette |
| Bacillota | *Lactococcus lactis* | GC984 | THY | 37 | Gift from Michael Surette |
| Bacillota | *Lactococcus lactis* | GC993 | THY | 37 | Gift from Michael Surette |
| Bacillota | *Listeria monocytogenes* | 10403S | BHI | 37 | (11) |
| Bacillota | *Listeria monocytogenes* | EGD-e | BHI | 37 | (12) |
| Bacillota | *Listeria monocytogenes* | CFSAN06B43 | BHI | 37 | |
| Bacillota | *Listeria swaminathanaii* | FSL L7.0020 | BHI | 30 | |
| Bacillota | *Staphylococcus aureus* | HG003 | TSB | 37 | (13) |
| Bacillota | *Staphylococcus aureus* | Newman | TSB | 37 | (14) |
| Bacillota | *Staphylococcus aureus* | USA300 | TSB | 37 | (15) |
| Bacillota | *Staphylococcus aureus* | RN4220 | TSB | 37 | (16) |
| Bacillota | *Staphylococcus aureus* | WCC 0119 | TSB | 37 | Gift from Gerard Wright |
| Bacillota | *Staphylococcus epidermidis* | 14.1.R1 | TSB | 37 | (17) |
| Bacillota | *Streptococcus intermedius* | B196 | THY | 37 | (18) |
| Bacillota | *Streptococcus pneumoniae* | ATCC 6303 | THY | 37 | (19) |
| Bacillota | *Streptococcus pyogenes* | MGAS5005 | THY | 37 | (20) |
| Bacteriodota | *Flavobacterium johnsoniae* | UW101 | LB | 30 | Gift from Mark McBride |
| Pseudomonadota | *Acinetobacter baylyi* | ADP1 | BHI | 37 | (21) |
| Pseudomonadota | *Burkholderia cenocepacia* | K56-2 | LB | 37 | (22) |
| Pseudomonadota | *Burkholderia kururiensis* | | LB | 37 | Gift from Joseph Mougous |
| Pseudomonadota | *Burkholderia vietnamiensis* | G4 | LB | 37 | (23) |
| Pseudomonadota | *Citrobacter freundii* | ATCC 8090 | LB | 37 | (24) |
| Pseudomonadota | *Enterobacter cloacae* | ATCC 13047 | LB | 37 | (25) |
| Pseudomonadota | *Escherichia coli* | MG1655 | LB | 37 | (26) |
| Pseudomonadota | *Escherichia coli* | Nissle 1917 | LB | 37 | (27) |
| Pseudomonadota | *Escherichia fergusonii* | ATCC 35469 | LB | 37 | (28) |
| Pseudomonadota | *Edwardsiella tarda* | ATCC 15947 | LB | 37 | (29) |

| | | | | | |
|---|---|---|---|---|---|
| Pseudomonadota | *Pseudomonas aeruginosa* | PAO1 | LB | 37 | (30) |
| Pseudomonadota | *Pseudomonas putida* | KT2440 | LB | 30 | (31) |
| Pseudomonadota | *Pseudomonas savastanoi* | pv. phaseolicola 1448a | LB | 30 | (32) |
| Pseudomonadota | *Ralstonia picketii* | ATCC 27511 | LB | 37 | (33) |
| Pseudomonadota | *Shigella flexneri* | 2a str. 301 | LB | 37 | Gift from Brian Coombes |
| Pseudomonadota | *Serratia proteomaculans* | 568 | LB | 37 | (34) |
| Pseudomonadota | *Vibrio cholerae* | El Tor N16961 | LB | 37 | (35) |
| Pseudomonadota | *Vibrio parahaemolyticus* | RIMD2210633 | LB | 37 | (36) |
| Pseudomonadota | *Xenorhabdus nematophilus* | AN6 | LB | 37 | (37) |

**Table S2:** Strains used in this study

| Organism | Genotype | Description | Reference |
|---|---|---|---|
| *E. coli* XL-1 Blue | *recA1 endA1 gyrA96 thi-1 hsdR17 supE44 relA1 lac* [F′ *proAB lacI*$^{q}$ ZΔ*M15* Tn*10* (Tet$^{R}$)] | Cloning strain | Novagen |
| *E. coli* DC10b | mcrA Δ(mrr-hsdRMS-mcrBC) φ80lacZΔM15 ΔlacX74 recA1 araD139 Δ(ara-leu)7697 galU galK rpsL endA1 nupG Δdcm | Methylase-deficient universal cloning strain for *E. coli/S. aureus* shuttle vectors | (38) |
| *E. coli* BL21 (DE3) pLysS | F- ompT gal dcm lon hsdSB(rB$^{-}$ mB$^{-}$) λ(DE3) pLysS(cm$^{R}$) | Protein expression strain | Novagen |
| *Staphylococcus aureus* USA300 | Wild-type | *S. aureus* ABP system expression strain | (15) |
| *Staphylococcus aureus* HG003 | Wild-type | AbpT-susceptible *S. aureus* strain | (13) |
| *Staphylococcus aureus* WCC 0119 | Wild-type | *S. aureus* human clinical isolate | Gift from Gerard Wright |
| *Staphylococcus aureus* MRSA252 | Wild-type | *S. aureus* clinical isolate | (39) |
| *Bacillus subtilis* PY79 | Wild-type | AbpT-susceptible *B. subtilis* strain | (4) |
| *Lactococcus lactis* GC982 | Wild-type | AbpT-susceptible *L. lactis* strain | Gift from Michael Surette |
| *Listeria monocytogenes* 10403S | Wild-type | AbpT-susceptible *L. monocytogenes* strain | (11) |
| *Corynebacterium glutamicum* MB001 | Wild-type | AbpT-susceptible *C. glutamicum* strain | (1) |

**Table S3:** Plasmids used in this study

| Plasmid | Relevant features | Reference |
|---|---|---|
| pBAD33 | Expression vector with *araBAD, ara* promoter, ChlorR | (40) |
| pBAD33::BC_3239_S28_VSVG | AbpT$^{Bc}$ expression vector | This study |
| pET29b | Expression vector with *lacI*, T7 promoter, C-terminal His6 tag, KanR | Novagen |
| pET29b::CCAS_05590_S46-CT_His6 | AbpP$^{Cc}$ expression vector | This study |
| pET29b::CCAS_05590_S46-CT_S210A_His6 | AbpP$^{Cc}_{S210A}$ expression vector | This study |
| pET29b::FC756_12635_S21-CT_His6 | AbpP$^{Lm}$ expression vector | This study |
| pETDuet-1 | Co-expression vector with *lacI*, T7 promoter, N-terminal His6 tag in MCS-1, AmpR | Novagen |
| pETDuet::His6_BG04_5990_S22-CT_H211A | AbpT$^{Pm}_{H211A}$ expression vector | This study |
| pETDuet::His6_BG04_5990_S22-CT::BG04_5991 | AbpT$^{Pm}$/AbpI$^{Pm}$ co-expression vector | This study |
| pETDuet::His6_CCAS_05600_E55-CT_H199A | AbpT$^{Cc}_{H205A}$ expression vector | This study |
| pETDuet::His6_CCAS_05600_E55-CT_H205A | AbpT$^{Cc}_{H211A}$ expression vector | This study |
| pETDuet::His6_CCAS_05600_E55-CT::CCAS_05595 | AbpT$^{Cc}$/AbpI$^{Cc}$ co-expression vector | This study |
| pETDuet::His6_FC756_12640_N29-CT_H211A | AbpT$^{Lm}_{H211A}$ expression vector | This study |
| pPSV39-CV | Expression vector with *lacI*, *lacUV5* promoter, GmR | (41) |
| pPSV39-CV::BC_3238_FLAG | AbpI$^{Bc}$ expression vector | This study |
| pPSV39-CV::BG04_5991 | AbpI$^{Pm}$ expression vector | This study |
| pPSV39-CV::SLD11_001950_FLAG | AbpI$^{Sa}$ expression vector | This study |
| pPSV39-CV::LYSIN_01453_FLAG | AbpI$^{Ls}$ expression vector | This study |
| pPSV39-CV::FB379_14220_FLAG | AbpI$^{Ac}$ expression vector | This study |
| pRAB11 | *E. coli/S. aureus* shuttle vector with *tetR*, P*tet* promoter, cmlR, ampR | (42) |
| pRAB11::SLD11_001950-001952 | ABP$^{Sa}$ system *S. aureus* expression vector | This study |

| | | |
|---|---|---|
| pRAB11::SLD11_001950-001952_001951_FLAG | ABP$^{Sa}$ system *S. aureus* expression vector encoding AbpT$^{Sa}$-FLAG | This study |
| pRAB11::SLD11_001950-001952_001951_H199A | ABP$^{Sa}$ system *S. aureus* expression vector encoding AbpT$^{Sa}_{H199A}$ | This study |
| pRAB11::SLD11_001950-001952_001951_H205A | ABP$^{Sa}$ system *S. aureus* expression vector encoding AbpT$^{Sa}_{H205A}$ | This study |
| pRAB11::SLD11_001950-001952_001952_S205A_001951_FLAG | ABP$^{Sa}$ system *S. aureus* expression vector encoding AbpT$^{Sa}$-FLAG, AbpP$^{Sa}_{S205A}$ | This study |
| pSCrhaB2-CV | Expression vector with P*rhaB*, C-terminal VSV-G tag, TmpR | (43) |
| pSCrhaB2-CV::BG04_5990_S22-CT_H211A_VSVG | AbpT$^{Pm}_{H211A}$ expression vector | This study |
| pSCrhaB2-CV::BG04_5990_S22-CT_H215A_VSVG | AbpT$^{Pm}_{H215A}$ expression vector | This study |
| pSCrhaB2-CV::BG04_5990_S22-CT_VSVG | AbpT$^{Pm}$ expression vector | This study |
| pSCrhaB2-CV::CCAS_05600_E55-CT_VSVG | AbpT$^{Cc}$ expression vector | This study |
| pSCrhaB2-CV::SAR2788_26-CT_K222A_VSVG | AbpT$^{Sa2}_{K222A}$ expression vector | This study |
| pSCrhaB2-CV::SAR2788_26-CT_VSVG | AbpT$^{Sa2}$ expression vector | This study |
| pSCrhaB2-CV::SLD11_001951_S33-CT_H199A_VSVG | AbpT$^{Sa}_{H199A}$ expression vector | This study |
| pSCrhaB2-CV::SLD11_001951_S33-CT_H205A_VSVG | AbpT$^{Sa}_{H205A}$ expression vector | This study |
| pSCrhaB2-CV::SLD11_001951_S33-CT_VSVG | AbpT$^{Sa}$ expression vector | This study |
| pSCrhaB2-CV::LYSIN_01454_S24-CT_VSVG | AbpT$^{Ls}$ expression vector | This study |
| pSCrhaB2-CV::LYSIN_01454_S24-CT_H226A_VSVG | AbpT$^{Ls}_{H226A}$ expression vector | This study |
| pSCrhaB2-CV::LYSIN_01454_S24-CT_H258A_VSVG | AbpT$^{Ls}_{H226A}$ expression vector | This study |
| pSCrhaB2-CV::FB379_14221_S30-CT_VSVG | AbpT$^{Ac}$ expression vector | This study |
| pSCrhaB2-CV::FB379_14221_S30-CT_E213Q_VSVG | AbpT$^{Ac}_{E213Q}$ expression vector | This study |
| pSCrhaB2-CV::FB379_14221_S30-CT_K215A_VSVG | AbpT$^{Ac}_{Q215A}$ expression vector | This study |
| pSCrhaB2-CV::FB379_14221_S30-CT_Q228A_VSVG | AbpT$^{Ac}_{Q228A}$ expression vector | This study |
